# Cysimiditides: RiPPs with a Zn-tetracysteine motif and aspartimidylation

**DOI:** 10.1101/2024.10.02.616296

**Authors:** Angela Zhu, Li Cao, Truc Do, A. James Link

**Affiliations:** Department of Chemical and Biological Engineering, Princeton University, Princeton, NJ 08544, United States; Department of Chemistry, Princeton University, Princeton, NJ 08544, United States; Department of Molecular Biology, Princeton University, Princeton, NJ 08544, United States

## Abstract

Aspartimidylation is a post-translational modification found in multiple families of ribosomally synthesized and post-translationally modified peptides (RiPPs). We recently reported on the imiditides, a new RiPP family in which aspartimidylation is the class-defining modification. Imiditide biosynthetic gene clusters encode a precursor protein and a methyltransferase that methylates a specific Asp residue, converting it to aspartimide. A subset of imiditides harbor a tetracysteine motif, so we have named these molecules cysimiditides. Here, using genome mining we show that there are 56 putative cysimiditides predicted in publicly available genome sequences, all within actinomycetota. We successfully heterologously expressed two examples of cysimiditides and showed that the major products are aspartimidylated and that the tetracysteine motif is necessary for expression. Cysimiditides bind a Zn^2+^ ion, presumably at the tetracysteine motif. Using *in vitro* reconstitution of the aspartimidylation reaction, we show that Zn^2+^ is required for methylation and subsequent aspartimidylation of the precursor protein. An AlphaFold 3 model of the cysimiditide from *Thermobifida cellulosilytica* shows a hairpin structure anchored by the Zn^2+^-tetracysteine motif with the aspartimide site in the hairpin loop. An AlphaFold 3 model of this cysimiditide in complex with its cognate methyltransferase suggests that the methyltransferase recognizes the Zn^2+^-tetracysteine motif to correctly dock the precursor protein. Cysimiditides expand the set of experimentally-confirmed RiPPs harboring aspartimides, and represent the first RiPP class that has an obligate metal ion.

**Table of Contents Figure:** 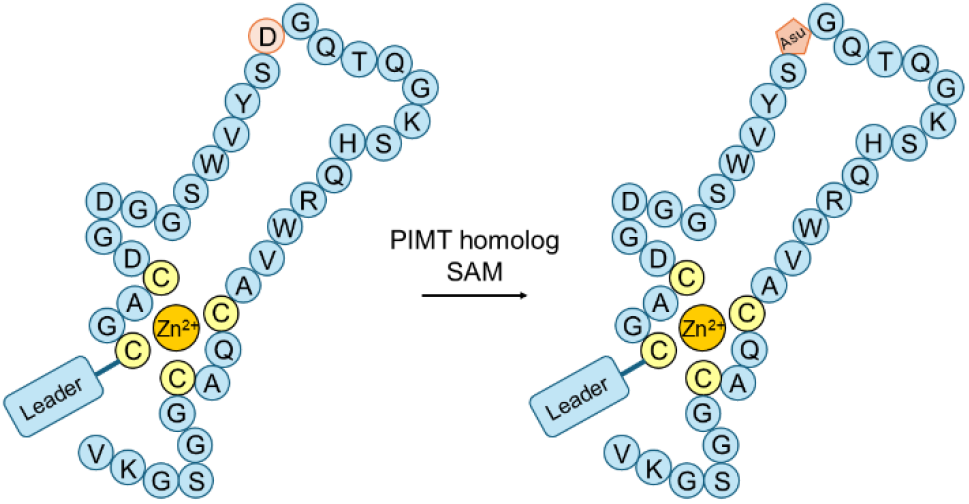

---

The aspartimide functional group, also known as aspartyl succinimide or cyclic imide, is a succinimide that introduces a rigidifying 5-membered ring into a peptide backbone. Aspartimides are found in proteins as intermediates in protein repair pathways^1-2^ and in intein-mediated protein splicing.^3-5^ Aspartimide moieties can also form from glycosylated aspartate residues.^6^ There are examples of unusually stable aspartimides within hyperthermophilic enzymes that help to either stabilize the fold of the enzyme^7-8^ or assist in substrate recognition.^9^ Another functional role for aspartimides (and 6-membered ring glutarimides) has been reported; peptides with these moieties at their C-terminus are recognized by the ubiquitin E3 ligase adapter cereblon and targeted for degradation.^10-11^ Recent work on the ribosomally synthesized and post-translationally modified peptide (RiPP) superfamily has revealed aspartimidylation as a modification that appears in multiple families of RiPPs including lanthipeptides, lasso peptides, and graspetides (Figure 1A).^12-17^ In all cases, the aspartimide is introduced after the class-defining modification(s). In contrast, we have recently reported on the imiditides, a class of RiPPs for which aspartimidylation is the class-defining modification.^18^ In these peptides, the aspartimide is installed on an unmodified linear peptide sequence.

**Figure 1:**
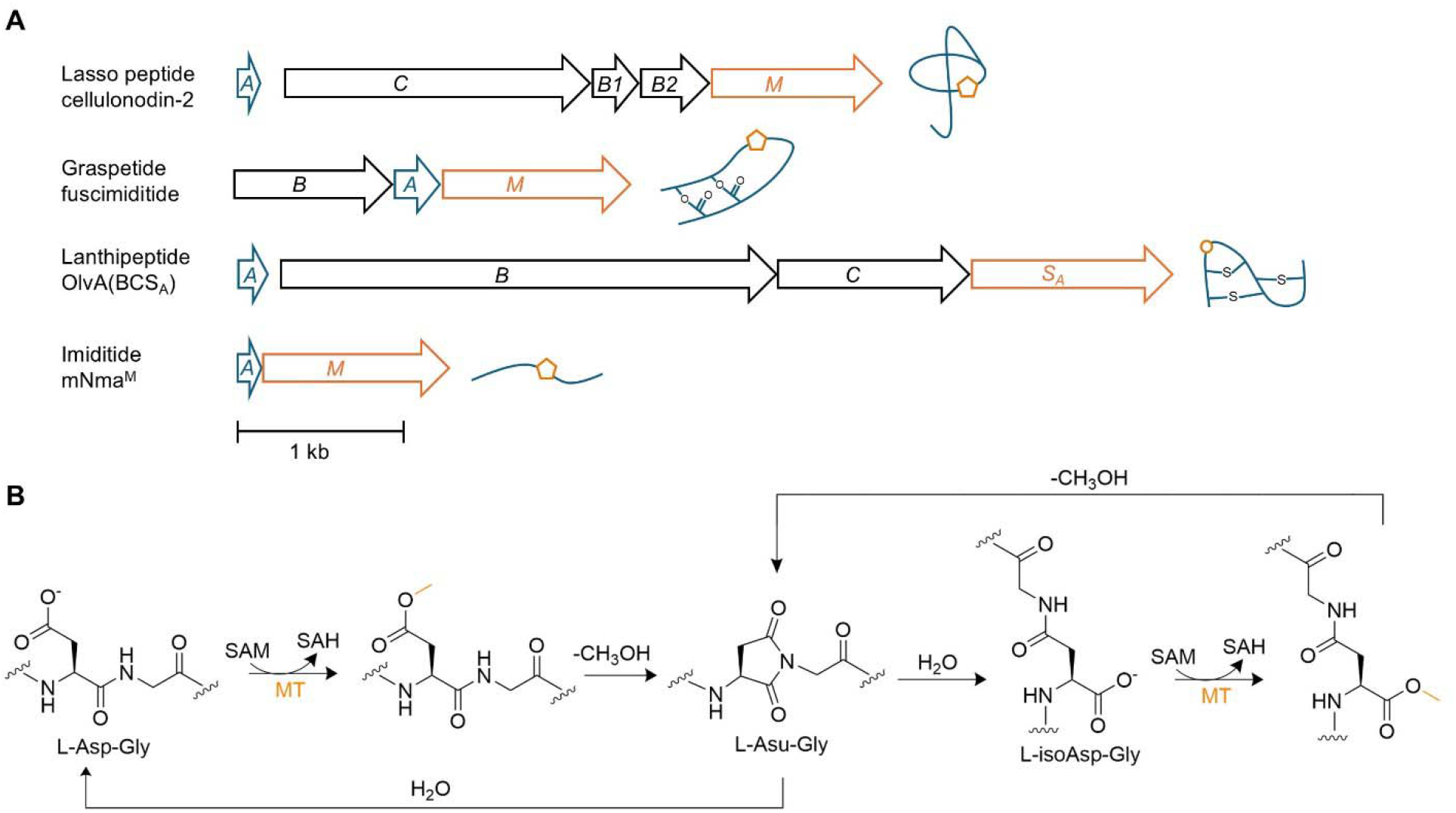
Aspartimidylation in ribosomally synthesized and post-translationally modified peptides (RiPPs). **(A)** Biosynthetic gene clusters (BGCs) across different RiPP families that contain a protein isoaspartyl methyltransferase (PIMT) homolog (orange). The PIMT homolog results in aspartimide modification (orange pentagons). In the lanthipeptide OlvA(BCS_A_), hydrolysis of the aspartimide leads to incorporation of isoAsp into the structure (orange circle). **(B)** Aspartimidylation pathway by PIMT homologs (MT) in RiPP BGCs. After methylation of the Asp sidechain, the adjacent backbone amide (Gly in this figure) attacks the methyl ester to form the aspartimide. The aspartimide can hydrolyze into isoAsp or back into Asp. The methyltransferase in the fuscimiditide BGC has been shown to also modify isoAsp residues. SAM is *S*-adenosyl methionine, and SAH is *S*-adenosyl homocysteine.

In our initial study, we found 670 putative imiditide biosynthetic gene clusters (BGCs) in public databases. These BGCs are comprised of only two genes encoding the precursor protein and a protein isoaspartyl methyltransferase (PIMT) homolog. The PIMT methylates a specific Asp residue within the precursor sequence, leading to the formation of the aspartimide moiety (Figure 1B). We noted that, among this set of imiditides, a small number of precursor sequences harbor conserved cysteine-containing motifs.^18^ This paper focuses on the biosynthesis of these Cys-containing imiditides, for which we suggest the name cysimiditides. Kim and coworkers have also published work on this class of molecules, suggesting the names “pamtide” for the imiditide family and type II pamtides for the cysimiditides.^17^ With a combination of genome mining approaches, we discover 56 putative cysimiditide BGCs and attempt heterologous expression of three different examples. We show that the ability of the cysimiditides to be recognized by the PIMT and aspartimidylated is strictly dependent on the presence of a Zn^2+^ ion coordinated by the conserved Cys residues.

## Results

### Discovery of cysimiditides

Our laboratory previously identified 670 putative imiditide biosynthetic gene clusters (BGCs) by using a C-terminal sequence motif conserved among imiditide methyltransferases as a bioinformatic handle for genome mining.^18^ We noticed that a subset of these imiditide sequences (19 out of 670) have two conserved CXXCXGXG amino acid motifs that flank the putatively aspartimidylated Asp residue (Figure S1). We used HHpred to identify homologous proteins that share these motifs.^19-20^ Namely, tetracysteine motifs are observed in zinc-binding proteins like chaperone protein DnaJ and inhibitor protein anti-*trp* RNA-binding attenuation protein (anti-TRAP) (Figure S2).^21-25^ We therefore decided to call this imiditide subset cysimiditides after the conserved cysteine residues.

We also used NmaM, the methyltransferase in a characterized imiditide BGC from *Nonomuraea maritima*,^18^ as the input for a sequence similarity network (SSN) of 1000 sequences (Figure S3).^26-27^ The SSN revealed two putative cysimiditide methyltransferases in thermophiles *Thermobifida cellulosilytica* and *Thermobifida fusca*. Our laboratory has previously characterized the aspartimidylated lasso peptide cellulonodin-2 and the aspartimidylated graspetide fuscimiditide from these organisms, respectively.^13-14^ Since these two cysimiditide methyltransferases did not appear in the results of our imiditide genome mining, we performed BLASTP searches on them and their associated cysimiditide precursors. We identified 10 additional putative cysimiditide BGCs from the 18 BLASTP results for the *T. cellulosilytica* cysimiditide precursor TceA and 25 more BGCs from the 100 BLASTP results for the associated methyltransferase TceM. BLASTP results for the *T. fusca* cysimiditide precursor TfuA and methyltransferase TfuM did not yield any cysimiditides not already found from the TceA and TceM results. Figure S4 summarizes the method of identifying the 56 total cysimiditide BGCs listed in Table S3.

The putative cysimiditide BGCs identified are all from actinobacterial genomes, isolated from both terrestrial and marine environments. Most are from the family *Nocardiopsaceae*, and the two most represented genera are *Nocardiopsis* and *Frankia* (Figure S5). The putatively modified Asp residues (as indicated by Clustal Omega alignment^28^) are all followed by Gly residues (Figure S1), as is commonly seen in aspartimidylated lanthipeptides and graspetides.^12, 14-15^ The number of residues between the CXXCXGXG motifs varies from 14 to 22, and the first cysteine motif sometimes varies as CXXCXG<u>XX</u>G or CXXCXG<u>XX</u>. Neighboring genes of cysimiditide BGCs were also examined using webFlaGs.^29^ Although no neighboring genes are universally conserved, 19 cysimiditide BGCs have helix-turn-helix transcriptional regulators nearby, and 15 of these regulators also have a DUF397 domain-containing protein nearby (Figure S6).^29^

### Heterologous expression of three cysimiditides

We chose the cysimiditides from *T. fusca* and *T. cellulosilytica* and a cysimiditide from *Nocardiopsis alba* for heterologous expression in *E. coli* (Figure 2A). The first two were chosen as their putative producing organisms are thermophilic, and our laboratory has successfully characterized RiPPs from these thermophilic organisms in the past.^13-14, 30^ The cysimiditide from *Nocardiopsis alba* was chosen because it is from a mesophilic organism, the *Nocardiopsis* genus was highly represented in our list of cysimiditides (Figure S5), and its precursor sequence varies sufficiently from the other two sequences. For heterologous expression, we added a N-terminal His_6_-tag to the precursor (*A* gene) and placed it under an IPTG-inducible T5 promoter, while we placed the untagged methyltransferase (*M* gene) on a separate plasmid under an IPTG-inducible T7 promoter. Expressing the precursors His_6_-TfuA and His_6_-NalA alone yielded species with observed monoisotopic masses of 6894.90 Da and 6733.94 Da, respectively, corresponding to the full proteins with cleavage of the N-terminal methionines (expected masses 6895.15 Da and 6733.96 Da, respectively) (Figure 2B). However, His_6_-TceA was not able to be expressed (Figure S7). The TceA sequence has CQA<u>G</u> for the second cysteine motif instead of the expected CQA<u>C</u> (Figure 2A). We expect this glycine to have originally been cysteine, as the fourth cysteine is conserved in all the other cysimiditide precursors; additionally, the exact TceA precursor sequence with CQAC is observed in the cysimiditide BGC of *Thermobifida alba* (Figure S1). We therefore substituted CQA<u>G</u> in TceA back into CQA<u>C</u> through a single base change, which improved expression and yielded a single species with observed monoisotopic mass of 6744.83 Da (expected mass 6744.90 Da, with cleavage of the N-terminal methionine) (Figure 2B). All subsequent TceA constructs thus contain this G to C amino acid substitution unless otherwise noted.

**Figure 2:**
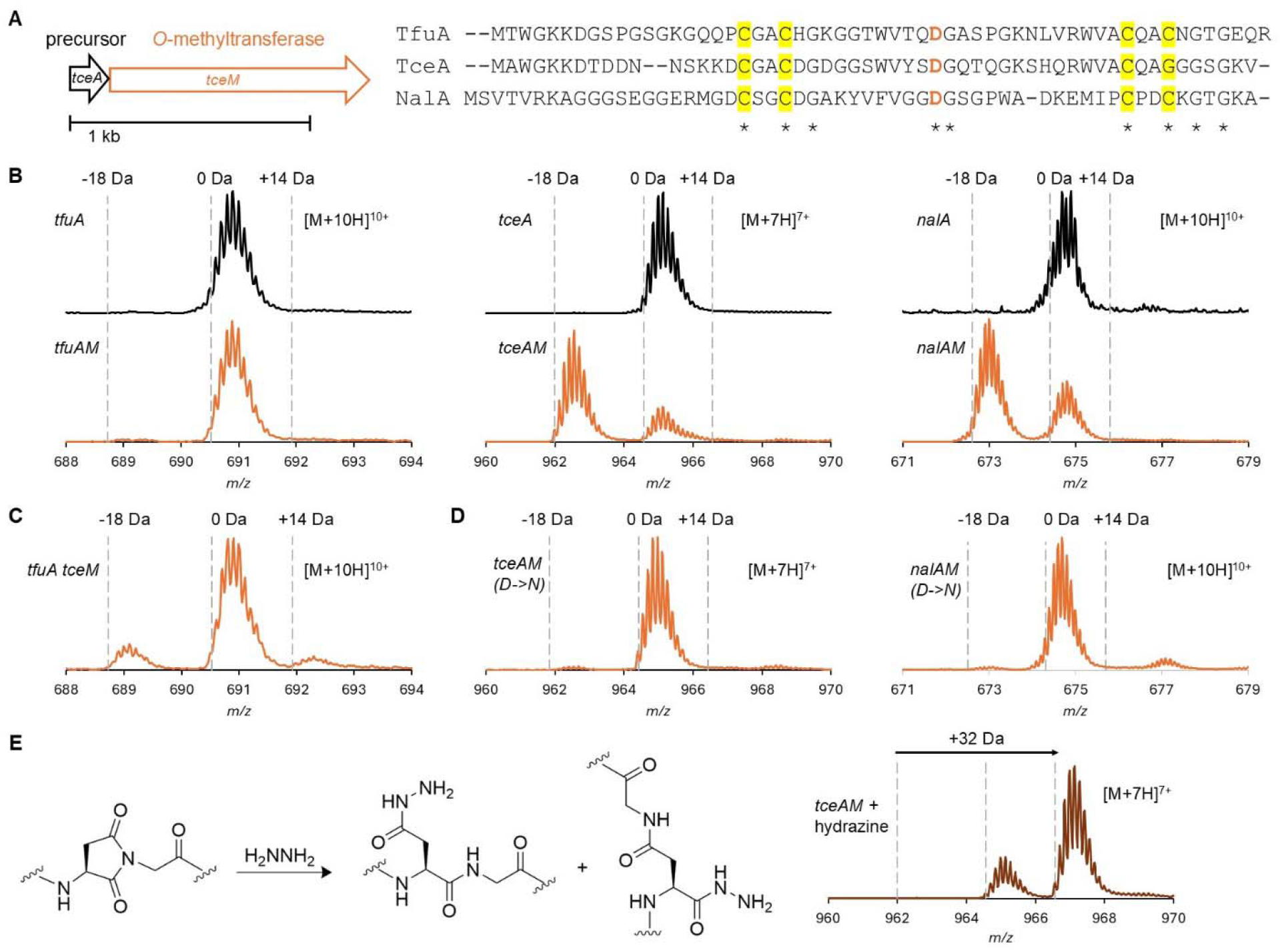
Heterologous expression of three cysimiditides in *E. coli* demonstrates that they are aspartimidylated. **(A)** Cysimiditide BGCs contain the *A* gene (precursor) and *M* gene (*O*-methyltransferase that is a PIMT homolog). The precursors contain two conserved CXXCXGXG motifs that flank the putative aspartimidylation site (colored orange). The precursor from *Thermobifida cellulosilytica* (TceA) contains a glycine in the fourth cysteine position. **(B)** Co-expression of the precursors with their associated methyltransferases resulted in the appearance of putatively aspartimidylated species (−18 Da), except for TfuM, which did not appear to modify His_6_-TfuA. The TceA sequence has been modified such that the second cysteine motif is CQA<u>C</u>. **(C)** His_6_-TfuA can be modified by TceM, suggesting that the TfuM enzyme is non-functional. **(D)** The aspartimide was installed on the conserved Asp between the cysteine motifs, as mutating this Asp to Asn in His_6_-TceA and His_6_-NalA abolished modification by their methyltransferases. **(E)** The reaction of aspartimide with hydrazine is expected to give a +32 Da increase. This mass shift was seen when reacting 2 M hydrazine with 10 μM of the co-expression product of His_6_-TceA and TceM for 30 minutes at room temperature.

Co-expression of the precursors with their associated methyltransferases resulted in the formation of -18 Da species as the major products for His_6_-TceA and His_6_-NalA, corresponding to putatively aspartimidylated species (Figure 2B). Surprisingly, His_6_-TfuA did not show dehydration after co-expression with TfuM (Figure 2B). However, His_6_-TfuA could instead be modified by co-expression with TceM (Figure 2C), which shares 68% sequence identity with TfuM (Figure S8). We thus conclude that TfuA is a functional substrate for methylation and aspartimidylation, but that the TfuM enzyme is defective in some way. We tried to restore function to TfuM by generating a series of TfuM/TceM chimeras, but these methyltransferase hybrids were still unable to modify His_6_-TfuA (Figure S9). His_6_-NalA was unable to be modified by TceM, most likely due to greater sequence differences between these two BGCs (Figure S10).

Next, we set out to confirm that the site of modification is the Asp in the conserved DG site between the cysteine motifs, as suggested by sequence alignment (Figure 2A). Altering this conserved Asp residue in His_6_-TceA and His_6_-NalA to Asn eliminated modification as expected (Figure 2D). Although NalA has a second Asp residue between the cysteine motifs, changing this residue to Asn did not eliminate modification (Figure S11). NalA also differs from TceA and TfuA in that it has a possible later start codon, located 3 amino acids before the first CXXC motif (Figure 2A). However, the construct we made with this later start codon did not express, while the construct with the earlier start codon did (Figure S12).

Since TceA and NalA were similarly modified, we focused on TceA for the rest of our studies. To confirm that the -18 Da species was aspartimidylated, we reacted 10 μM of the co-expression product of His_6_-TceA and TceM (Figure 2B) with 2 M hydrazine in 50 mM Tris-HCl at pH ∼7 for 30 minutes at room temperature (Figure 2E). The hydrazine reaction product showed the expected conversion of the -18 Da species into a +14 Da species, confirming the presence of aspartimide.^31^ We denote the aspartimidylated species as mTceA^M^, where m stands for modified and ^M^ represents modification by the methyltransferase. We also expressed a His_6_-SUMO-TceA construct with an observed average mass of 17891.56 Da (expected mass 17891.78 Da, with cleavage of the N-terminal methionine), which was likewise able to be modified by TceM, albeit less than His_6_-TceA was (Figure 3A).

**Figure 3:**
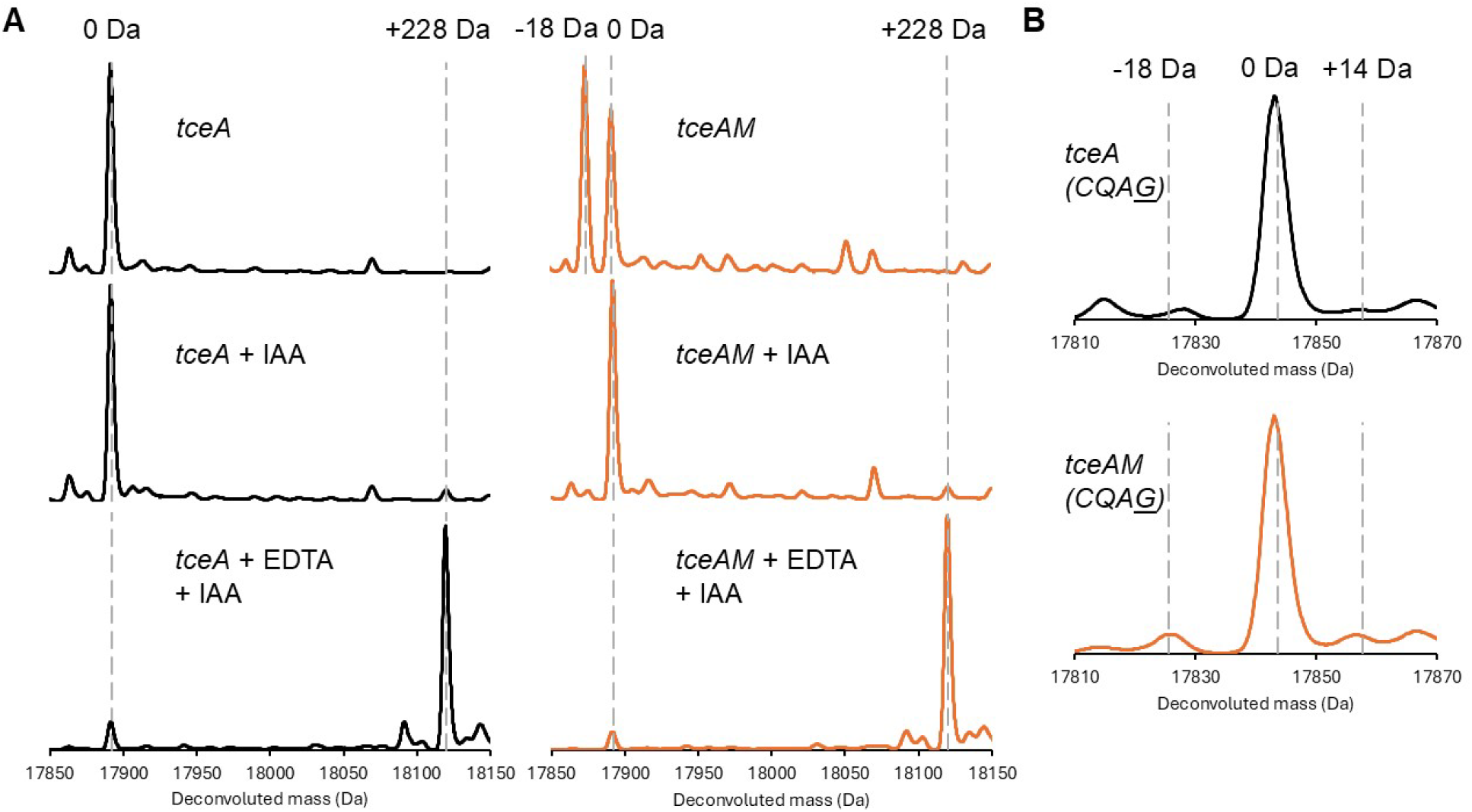
The cysteine motifs of TceA are necessary for aspartimidylation. **(A)** Alkylation reactions with iodoacetamide (IAA) of natively purified His_6_-SUMO-TceA and its co-expression product with TceM. Pre-treatment of the protein with 1 mM EDTA for 15 min at 37 °C is necessary for significant alkylation to occur. Prior to alkylation, the protein was reduced with TCEP at 55 °C, which also hydrolyzed the aspartimidylated species in the co-expression product. **(B)** His_6_-SUMO-TceA cannot be modified by TceM *in vivo* when its fourth cysteine is replaced by glycine.

### Zinc is required for cysimiditide modification

CXXCXGXG motifs are known to bind zinc in proteins such as DnaJ and anti-TRAP.^23, 25^ Zinc binding can contribute to protein stability; mutating the cysteines in anti-TRAP has been shown to decrease expression, most likely due to degradation of the mutants in the absence of zinc binding.^24^ This can explain our inability to express His_6_-TceA with glycine in the fourth cysteine position (Figure S7). We found that His_6_-SUMO-TceA (with all four cysteines) and its co-expression product with TceM were both unable to be alkylated with iodoacetamide unless first pre-treated with 1 mM EDTA for 15 min at 37 °C. This result implies that the cysteines in TceA are already occupied with zinc binding (Figure 3A).

Next, we used the zinc-binding reagent 4-(2-pyridylazo)resorcinol (PAR) to detect the presence of zinc in natively purified His_6_-TceA and His_6_-SUMO-TceA, their co-expression products with TceM, and His_6_-TceM. His_6_-TceM was included as a negative control since it was not expected to have zinc bound. As positive controls, we assessed the presence of zinc in zinc-binding proteins anti-TRAP from *Bacillus subtilis* and the cysteine-rich region of DnaJ from *Thermobifida cellulosilytica*, both of which we heterologously expressed in *E. coli* with His_6_-tags and natively purified (Figure S13). The binding of PAR to Zn^2+^ causes an increase in absorbance at 500 nm compared to the absorbance of PAR alone, which can be monitored to measure zinc binding (Figure S14).^32-33^ We first incubated the proteins with 5 M urea at 95 °C for 30 min to denature them and facilitate zinc release before mixing with a final concentration of 100 μM PAR. Greater zinc to protein ratios were observed in the TceA samples compared to the His_6_-TceM sample (Figure S14). Although we were not able to observe 1:1 zinc to protein ratios as expected, we attribute this to tight protein-zinc binding that PAR is unable to outcompete even with the denaturing condition. Indeed, previous quantifications of zinc binding to anti-TRAP and DnaJ have used strong thiol-binding reagents such as *p*-hydroxymercuribenzoate (HMB) or *p*-hydroxymercuriphenylsulfonic acid (PMPS) to more completely release zinc.^22, 24^

Finally, we were interested in seeing if zinc binding affects modification by TceM. We were able to express a His_6_-SUMO-TceA construct with glycine in the fourth cysteine position; the observed mass was 17843.38 Da compared to the expected mass of 17845.69 Da, indicating formation of one disulfide bond. TceM was unable to significantly modify this His_6_-SUMO-TceA glycine construct when co-expressed *in vivo* (Figure 3B). Collectively, these data suggest that TceA binds zinc *in vivo* and that zinc binding likely stabilizes the TceA structure for recognition by TceM.

### *In vitro* reconstitution of TceA aspartimidylation

Having established that multiple cysimiditides are aspartimidylated *in vivo*, we turned our attention to *in vitro* reconstitution of the aspartimidylation reaction. We performed a time course of His_6_-TceA modification by His_6_-TceM *in vitro* in 50 mM Tris-HCl pH 7.4, 400 μM SAM, 1 mM DTT, and 100 μM ZnCl_2_. Methylated species (+14 Da) appeared within 5 minutes (Figure S15), and by 3 h, the level of aspartimidylated species (−18 Da) was slightly more than that of the unmodified species (Figure 4A). Allowing the reaction to proceed for 6 or 16 h did not considerably increase the level of aspartimidylated species (Figure S16). Performing the reactions without adding 100 μM ZnCl_2_ (Figure 4B) yielded a similar amount of aspartimidylation between 1-3 h as when zinc was added, which agrees with the previously described results indicating that natively purified TceA carries its own zinc. Pre-incubating His_6_-TceA with 1 mM EDTA for 15 min at 37 °C before adding the other reaction components blocked aspartimidylation from occurring, showing directly that zinc was necessary for this modification (Figure 4C). *In vitro* reactions were also performed with His_6_-TceA that was urea purified, incubated with 3 mM EDTA for 15 min at 37 °C, and then desalted to remove both zinc and EDTA. These reactions showed only a low level of aspartimidylated species (Figure 4D), while re-adding 100 μM ZnCl_2_ restored high levels of modification (Figure 4E), further demonstrating the necessity of zinc. In contrast, adding 100 μM of iron(II) sulfate heptahydrate or nickel(II) sulfate hexahydrate did not restore the high level of modification (Figure S17).

**Figure 4:**
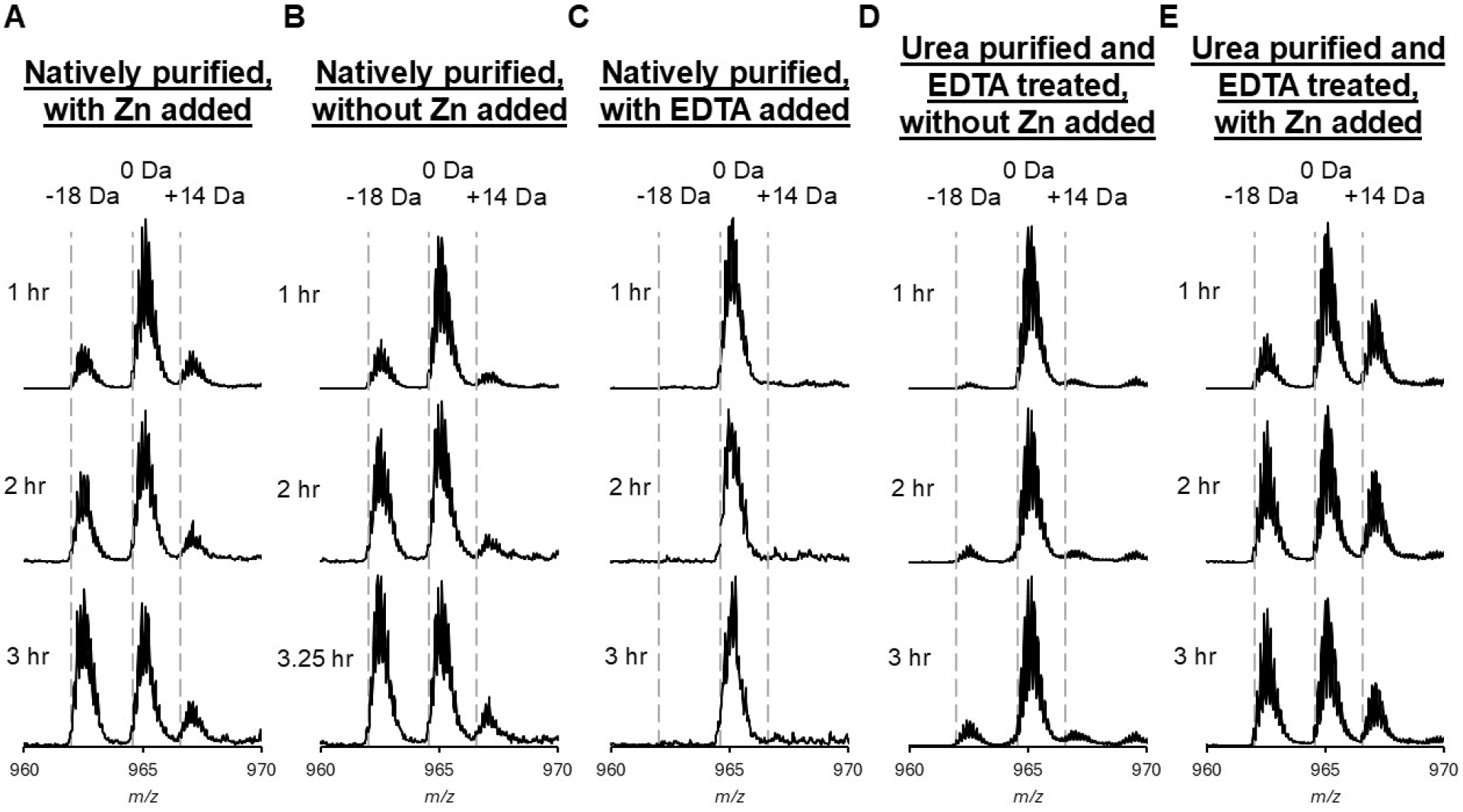
*In vitro* reactions of His_6_-TceA. **(A)** Natively purified His_6_-TceA (5 μM) was reacted with 1 μM His_6_-TceM in 50 mM Tris-HCl, pH 7.4, 400 μM SAM, 1 mM DTT, and 100 μM ZnCl_2_. Reactions were incubated at 37 °C for 1-3 h before quenching with formic acid and analysis using LC-MS. The methylated species (+14 Da) was present throughout, and the amount of aspartimidylated species (−18 Da) increased over time. **(B)** Reactions were performed in the same conditions as (A) but without adding 100 μM ZnCl_2_. The amount of modification was similar to that seen in (A). **(C)** Reactions were performed in the same condition as (B), but His_6_-TceA was pre-incubated with 1 mM EDTA for 15 min at 37 °C before the other reaction components were added. This abolished modification. **(D)** Urea purified elutions of His_6_-TceA were incubated with EDTA and run through a PD-10 desalting column to remove bound zinc and EDTA. Reactions with this substrate were performed in the same conditions as (B). The amount of aspartimidylated species seen was much lower. **(E)** Reactions were performed with the same substrate as (D) in 100 μM ZnCl_2_. High levels of aspartimidylated species were restored with the addition of zinc.

In all of our heterologous expression and *in vitro* experiments with TceA, we never found conditions in which TceA was fully converted to the aspartimidylated form. This result contrasts with what is observed with aspartimidylated lasso peptides and graspetides where nearly all of the peptide was converted to the aspartimide form.^13-15^ This led us to the hypothesis that the aspartimide in mTceA^M^ is more labile than that observed in aspartimidylated lasso peptides and graspetides. We incubated the His_6_-TceA and TceM co-expression product (Figure 2B) at room temperature at pH 7.4 for four hours and analyzed a sample every hour using LC-MS. This brief time at near-neutral pH was enough for partial hydrolysis of the aspartimide to occur (Figure S18). We performed a 16 h trypsin digest of His_6_-TceA and its TceM co-expression product, during which the aspartimidylated species mTceA^M^ hydrolyzed completely. mTceA^M^ hydrolyzed into a mix of isoAsp and Asp, as shown by two trypsin fragments with the same mass of 2204.78 Da (expected mass 2204.84 Da) but different retention times on LC-MS (Figure 5A). The presence of isoAsp was further supported by successful reaction of the hydrolyzed co-expression product with human PIMT, which is known to only methylate and dehydrate isoAsp (Figure 5B). His_6_-TceM was also able to re-modify the hydrolyzed product (Figure 5C).

**Figure 5:**
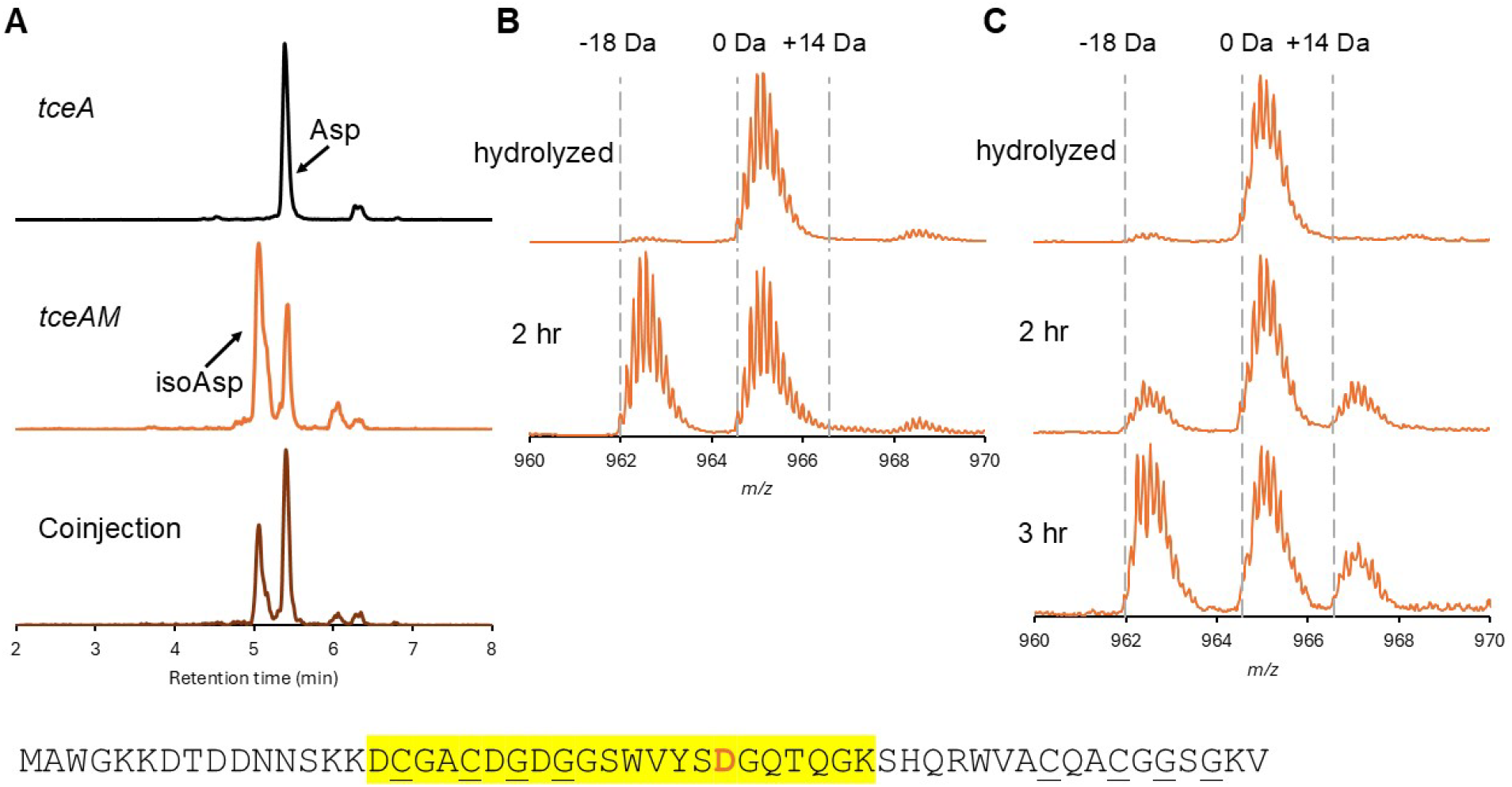
Hydrolysis of mTceA^M^. **(A)** His_6_-TceA and its co-expression product with TceM were digested with trypsin for 16 h at 37 °C in 50 mM Tris-HCl pH 8 and analyzed on LC-MS. The trypsin digest of the co-expression product showed two peaks with different retention times corresponding to isoAsp- and Asp-containing fragments. The sequence of the trypsin fragment is highlighted below. **(B)** *In vitro* reaction of 5 μM of hydrolyzed co-expression product with 1 μM human PIMT in 50 mM Tris-HCl, pH 7.4, 400 μM SAM, 1 mM DTT, and 100 μM ZnCl_2_ at 37 °C. The co-expression product was hydrolyzed for 16 h at room temperature in 50 mM Tris-HCl, pH 8. Human PIMT modified the hydrolyzed product, confirming the presence of isoAsp after aspartimide hydrolysis. **(C)** *In vitro* reactions of 5 μM of hydrolyzed co-expression product with 1 μM His_6_-TceM in the same conditions as (B). TceM can re-modify the hydrolyzed product.

## Discussion

Here we have described the cysimiditides, a RiPP subfamily characterized by the presence of an aspartimide modification and a zinc ion coordinated by 4 Cys residues. The cysimiditides are a subdivision of the larger imiditide class; there are several hundred imiditides in published genomes, and 56 of them are cysimiditides. The cysimiditides are found exclusively in actinobacteria/actinomycetota. Cysimiditides represent another example of a RiPP natural product that is modified with the aspartimide moiety. Both lasso peptides and graspetides with aspartimides have been isolated from *E. coli* in heterologous expression studies.^13-16^ In contrast to our findings, Kim and colleagues report little aspartimidylated product in their heterologous expression of cysimiditides (named type II pamtides in their work).^17^ We believe that this discrepancy is due to differences in how the cysimiditide was processed after *E. coli* expression. While in our experiments we immediately buffer-exchanged the purified cysimiditide into a buffer at pH 7.4, Kim and colleagues added a TEV protease digestion step at pH 8 for 16 h. Under these conditions we expect that any aspartimide that may be present would be hydrolyzed. Our interpretation of our data here and in other publications is that the intended final RiPP product *in cellulo* is aspartimidylated.^13-14^ If the cysimiditide remains in the cell, it will retain the aspartimide modification, as long as SAM and methyltransferase are present. However, if the cysimiditide is exported into neutral or basic conditions, the aspartimide will hydrolyze into a mixture of Asp and isoAsp. If cysimiditides are exported into an acidic environment, such as the stomach of animal, the aspartimide will persist for longer periods of time. Studies in cysimiditide native producers are needed to understand the transport of these molecules.

This work on cysimiditides also illustrates that aspartimide moieties in different classes of RiPPs have differences in their hydrolytic stability. In the case of the cysimiditides, we never observed full conversion of the precursor to its aspartimidylated form, either in heterologous expression experiments or after 16 h *in vitro*. The accumulation of the aspartimide can be explained using a kinetic framework, as we have previously published for the aspartimidylated lasso peptide lihuanodin.^34^ Aspartimidylated lasso peptides and graspetides are fully converted to the aspartimide form because the rate of hydrolysis is small compared to the rate of methylation and dehydration. However, for the cysimiditides (and imiditides more broadly) we propose that the rate of hydrolysis competes with the rates of methylation and dehydration, leading to the mixture of products we observed.

The cysimiditides are also unique among RiPPs in that their structure requires the coordination of a metal ion. While other RiPPs, including methanobactin and related compounds^35-37^ and the lanthipeptide noursin^38^ have metal-binding functions, we believe that cysimiditides are the first example of a RiPP in which a metal ion is integral to the stability (Figure S7) and to the installation of the post-translational modification (Figure 4) on the RiPP. The AlphaFold 3 predicted structure of the *T. cellulosilytica* cysimiditide TceA is a hairpin structure (Figure 6A) that is anchored by the Zn^2+^ tetracysteine motif.^39^ A key consideration with RiPPs that are modified by this class of methyltransferase is substrate specificity. If the TceM methyltransferase were promiscuous and methylated Asp residues throughout the proteome, then extensive damage to the proteome would occur. Thus, TceM and other aspartimidylating methyltransferases must be highly specific for only the RiPP substrate. In the case of lanthipeptides, lasso peptides, and graspetides, this specificity is enforced by the fact that only modified peptides are substrates for their cognate methyltransferases.^12-15^ In the case of imiditides, substrate specificity is enforced by extensive charge-charge interactions.^18^ To understand the potential substrate specificity in cysimiditide methylation, we built an AlphaFold 3 model of TceA/zinc bound to TceM (Figure 6B).^39^ The structure of the complex seems plausible given that the Asp31 residue of TceA that is methylated and aspartimidylated is bound next to the putative SAM binding site of TceM. The structural model of the complex also shows contacts between the Zn^2+^ tetracysteine motif of TceA and the C-terminal portion of TceM (Figure 6B). We propose that this unique structural motif of TceA is responsible for its specific recognition by TceM.

**Figure 6:**
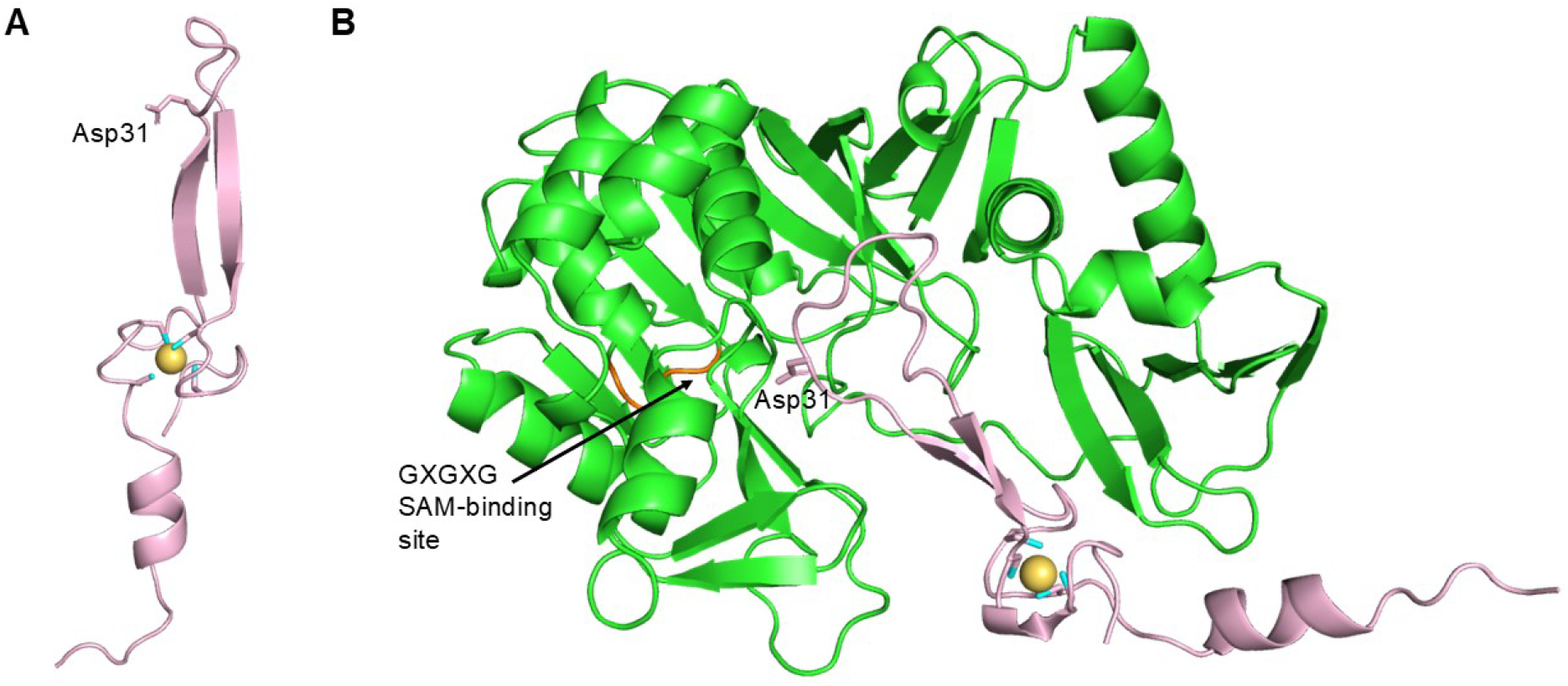
AlphaFold 3 models of TceA and TceM. **(A)** The AlphaFold 3 model of TceA is a hairpin structure with Asp31, the residue that is modified (in sticks), in the loop of the hairpin. The Zn^2+^ ion is shown in yellow, coordinated by the cysteines (sticks) in TceA. **(B)** When TceA and TceM are modeled as a complex, Asp31 (in sticks) of TceA (pink) is positioned near the GXGXG SAM-binding site (orange) of TceM (green). Residues near the cysteine motifs of TceA are also proximal to residues in the C-terminal domain of TceM.

### Supporting Information

Detailed materials and methods, additional mass spectrometry data, cysimiditide precursor and methyltransferase sequences and AlphaFold 3 structures, and 4-(2-pyridylazo)resorcinol (PAR) assay measurements.

### Accession Codes

The *Thermobifida cellulosilytica* and *Nocardiopsis alba* cysimiditides (TceM and NalM) are associated with the NCBI protein accession codes WP_083948071.1 and WP_234305832.1, respectively. The *Thermobifida fusca* cysimiditide methyltransferase (TfuM) is associated with the NCBI protein accession codes QOS59465.1 and WP_227480949.1. Anti-TRAP from *Bacillus subtilis* strain 168 and DnaJ from *Thermobifida cellulosilytica* are associated with the NCBI protein accession codes WP_003234807.1 and WP_068753762.1, respectively.

## Acknowledgments

This work was supported by NIH grant GM107036 to A.J.L. A.Z. was supported by an NSF Graduate Research Fellowship Program under Grant DGE-2039656. L.C. was supported by an NSF Graduate Research Fellowship Program under Grant DGE-1656466 and a Proctor Fellowship from Princeton University.

## For Table of Contents only

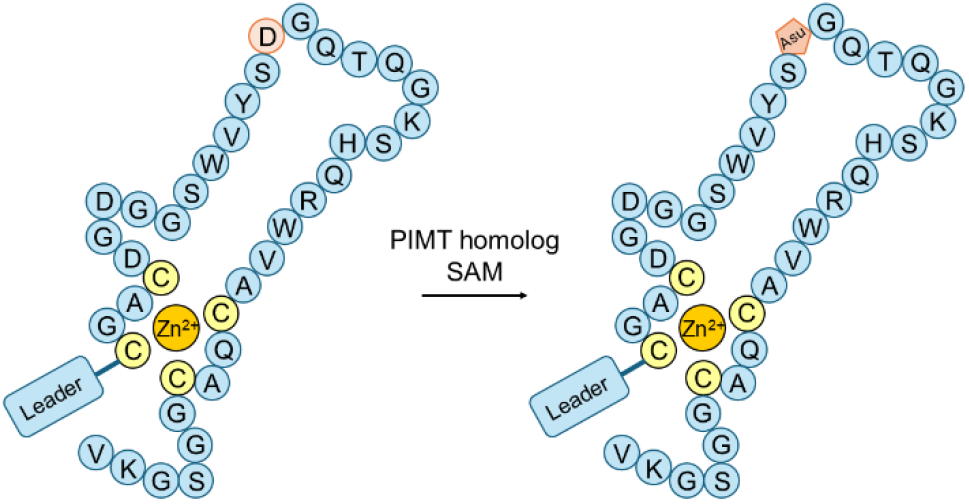

