## Supplemental Information for "Cysimiditides: RiPPs with a Zn-tetracysteine motif and aspartimidylation"

### Supplementary Materials and Methods

**Strains**

*Thermobifida fusca* (DSM 43792) and *Thermobifida cellulosilytica* (DSM 44535) were obtained from the German Collection of Microorganisms and Cell Cultures (DSMZ). *Nocardiopsis alba* strain TP-A0876 (NBRC 110039) was obtained from Japan’s National Institute of Technology and Evaluation (NITE) Biological Resource Center (NBRC). Organisms were grown according to suppliers’ recommended protocols, and genomic DNA was isolated using the Qiagen DNeasy Blood and Tissue Kit standard protocol.

**Cloning**

Plasmids were constructed through PCR amplification, restriction enzyme digestion, ligation, and transformation into chemically competent *E. coli* XL1-Blue on Lysogeny Broth (LB) plates supplemented with 100 ug/mL ampicillin or 50 ug/mL kanamycin as needed. Q5 DNA polymerase, restriction enzymes, antarctic phosphatase, and T4 DNA ligase were purchased from New England Biolabs (NEB), and all primers were purchased from Integrated DNA Technologies (IDT). Plasmids were extracted using Qigaen QIAprep Spin Miniprep Kit and sequenced using Sanger sequencing through Genewiz.

All primers used are listed in **Table S1**, and plasmid descriptions as well as the cloning methods for each plasmid are listed in **Table S2** at the end of the Methods section. To briefly describe the cloning of the *T. fusca* cysimiditide as a representative example, a region surrounding the *tfuAM* BGC was first PCR amplified from genomic DNA, digested using *Sac*I and *Hin*dIII, and ligated into pQE-80L to form pAZ1. *tfuM* was then PCR amplified using pAZ1 as a template, digested with *Bsa*I and *Hin*dIII, and ligated into the first multiple cloning site (MCS1) of *Nco*I- and *Hin*dIII-digested pRSF-Duet to form pAZ2. *tfuA* was PCR amplified using pAZ1 as a template, digested with *Bam*HI and *Hin*dIII, and ligated into pQE-80L to form pAZ12. The His_6_-tag in pAZ12 was also modified from MRGSHHHHHHGS to MSGSHHHHHHGS to eliminate N-terminal methylation. The mutation was introduced via primers, followed by digestion with *Eco*RI and *Hin*dIII and ligation. Similarly, *tceAM* and *nalAM* biosynthetic gene clusters were amplified from genomic DNA, digested using *Bam*HI and *Hin*dIII, and ligated into pAZ12 digested with *Bam*HI and *Hin*dIII. Using these plasmids as templates, the methyltransferases and precursors were then individually cloned into pRSF-Duet and pQE-80L vectors. The start codon of *nalA* was changed from GTG to ATG during this. Variants of TfuM, TceA, and NalA were cloned using the wild-type plasmids as templates and internal primers that introduced the relevant mutations, followed by overlap PCR, digestion, and ligation.

**Protein expression and purification**

Plasmids were transformed into *E. coli* BL21 (DE3) Δ*slyD* through electroporation. Cultures were grown overnight in LB media (supplemented with 100 ug/mL ampicillin and/or 50 ug/mL kanamycin as needed) shaking at 250 rpm at 37 ^o^C. The overnight culture was used to start a subculture at OD_600_ = 0.02 in 500 mL LB (supplemented with 100 ug/mL ampicillin and/or 50 ug/mL kanamycin as needed) shaking at 250 rpm at 37 ^o^C. Once the culture reached OD­­_600_ = 0.45-0.6, protein expression was induced with 1 mM IPTG, and the culture was moved to shaking at 250 rpm at room temperature (20-22 ^o^C) for 16-20 h. Cells were centrifuged at 4000 x *g* for 15 min at 4 ^o^C before proceeding to native or denaturing purification.

For native purification, all steps were performed on ice/at 4 ^o^C. The cell pellet (per 500 mL culture) was resuspended in 10 mL of 50 mM NaH_2_PO_4_, 300 mM NaCl, 10 mM imidazole, pH 8.0. The resuspension was incubated with 1 mg/mL lysozyme for 20 minutes and then sonicated for 12 cycles of 10 s on and 20 s off to lyse the cells. The lysate was centrifuged twice at 8000 x *g* for 15 minutes, with the supernatant decanted for additional centrifugation in between. The final supernatant was then mixed with 1 mL Ni-NTA resin (Qiagen) and incubated while rotating at 4 ^o^C for 1 h. The mixture was passed through an empty gravity column, followed by a 10 mL wash with 50 mM NaH_2_PO_4_, 300 mM NaCl, 20 mM imidazole, pH 8.0, and a second 10 mL wash with 50 mM NaH_2_PO_4_, 300 mM NaCl, 50 mM imidazole, pH 8.0. The protein was eluted with 50 mM NaH_2_PO_4_, 300 mM NaCl, 250 mM imidazole, pH 8.0, through eight 1 mL fractions. For proteins below 10 kDa, elution fractions 2-4 were pooled and buffer exchanged using a PD-10 desalting column (Bio-Rad) into 50 mM Tris-HCl, 100 mM NaCl, 10% glycerol, pH 7.4. For proteins above 10 kDa, elution fractions were pooled and buffer exchanged using 10 kDa or 30 kDa Amicon Ultra Centrifugal Filters as applicable into 50 mM Tris-HCl, 100 mM NaCl, 10% glycerol, pH 7.4. Human protein isoaspartyl methyltransferase (PIMT) was buffer exchanged into 50 mM Tris-HCl, 150 mM NaCl, 5 mM DTT, and 25% glycerol, pH 7.2.^1^ Purified proteins were stored in aliquots at -80 ^o^C.

For denaturing purification of His_6_-TceA, the cell pellet (per 500 mL culture) was resuspended in 10 mL of 100 mM NaH_2_PO_4_, 10 mM Tris, 8 M urea, pH 8.0. The resuspension was frozen at -80 ^o^C for 30 minutes, thawed in a water bath, and centrifuged twice at 8000 x *g* for 15 minutes. The final supernatant was then mixed with 1 mL Ni-NTA resin (Qiagen) and incubated while rotating at 4 ^o^C for 1 h. The mixture was passed through an empty gravity column, followed by a 10 mL wash with 100 mM NaH_2_PO_4_, 10 mM Tris, 8 M urea, pH 6.3, and a second 10 mL wash with 100 mM NaH_2_PO_4_, 10 mM Tris, 8 M urea, pH 5.9. The protein was eluted with 100 mM NaH_2_PO_4_, 10 mM Tris, 8 M urea, pH 4.5, through eight 1 mL fractions. Elution fractions 2-4 were pooled and buffer exchanged using a PD-10 desalting column (Bio-Rad) into 50 mM Tris-HCl, 100 mM NaCl, 10% glycerol, pH 7.4. If the removal of zinc was desired, elution fractions 2-4 were incubated with 3 mM EDTA for 15 min at 37 ^o^C before buffer exchange through the desalting column.

**Liquid chromatography-mass spectrometry (LC-MS)**

LC-MS analysis was performed using an Agilent 6530 QTOF connected to an Agilent 1260 LC system. Mass spectra were acquired using electrospray ionization (ESI) with the instrument in positive ion mode. The mobile phase A was water with 0.1% formic acid, and mobile phase B was acetonitrile with 0.1% formic acid. Intact proteins were run on a Xbridge Protein BEH C4 column (2.1 mm x 50 mm, 3.5 µM particle size, Waters). The following gradient was used for running His_6_-TceA on LC-MS due to its high polarity: 5% B from 0-1 minute, linear gradient from 5-50% B from 1-15 minutes, linear gradient from 50-90% B from 15-20 minutes, and 90% B for 5 minutes, with the first 0.5 min sent to waste. For running all other proteins, the following gradient was used: 10% B from 0-2 minute, linear gradient from 10-50% B from 2-15 minutes, linear gradient from 50-90% B from 15-20 minutes, and 90% B for 5 minutes, with the first 2 min sent to waste. Peptides from trypsin digestion were run on a Zorbax 300SB-C18 column (2.1 mm x 50 mm, 3.5 µM particle size, Agilent). The gradient used for the peptide column was: 10% B from 0-1 minute, linear gradient from 10-50% B from 1-20 minutes, linear gradient from 50-90% B from 20-25 minutes, and 90% B for 5 minutes, with the first 2 min sent to waste. Data were analyzed using Agilent MassHunter software. Deconvolution was performed using Agilent Bioconfirm software.

**Hydrazine reaction**

10 µM of the His_6_-TceA and TceM co-expression product was reacted with 2 M hydrazine (35% solution in water, Sigma-Aldrich) in 50 mM Tris-HCl in a total volume of 50 uL. HCl was added to the reaction such that the pH of the final reaction mixture was ~7, as indicated by a Fisherbrand pH indicator strip. The reaction was incubated at room temperature for 30 minutes and analyzed using LC-MS.

***In vitro* reactions**

*In vitro* reactions were performed using 5 µM TceA substrate, 1 µM of methyltransferase (His_6_-TceM or human PIMT), 400 µM SAM, 1 mM DTT (freshly made), and 100 µM ZnCl_2_ (if applicable) in 50 uL per reaction at 37 ^o^C. To hydrolyze the His_6_-TceA and TceM co-expression product, the protein was mixed with an equal volume of 50 mM Tris-HCl pH 8 and left for 16 h at room temperature. If EDTA was to be added to the reaction, then His_6_-TceA was pre-incubated with 1 mM EDTA for 15 min at 37 ^o^C before the other reaction components were added. *In vitro* reactions were quenched with 5 uL of 10% formic acid at the end of each time point and analyzed using LC-MS.

**Trypsin digestion**

His_6_-TceA or its co-expression product with TceM was digested with Sequencing Grade Modified Trypsin (Promega) in 1 mM DTT (freshly made) and 50 mM Tris-HCl pH 8 at 37 ^o^C for 16 h. A 1:100 (w/w) trypsin:protein ratio was used. The digestion was quenched with a final concentration of 1% formic acid and analyzed using LC-MS.

**Iodoacetamide reaction**

The relevant volume of purified His_6_-SUMO-TceA (or its co-expression product with TceM) correpsonding to 26.5 µg was mixed with 45 µL of 200 mM ammonium bicarbonate pH 8, and the total volume was then adjusted to 100 µL with sterile water. 5 µL of 200 mM TCEP-HCl in 200 mM ammonium bicarbonate pH 8 was added, and the reaction was incubated at 55 ^o^C for 1 h. 5 µL of 375 mM iodoacetamide in 200 mM ammonium bicarbonate pH 8 (freshly prepared and protected from light) was then added, followed by 30 min incubation at room temperature while protected from light. The reaction was analyzed using LC-MS.

If EDTA pretreatment was desired, 1 mM EDTA was added to the 100 uL mixture of protein, ammonium bicarbonate, and sterile water. The mixture was incubated at 37 ^o^C for 15 min. The rest of the reaction then proceeded identically as described above.

**Measurement of zinc binding**

200 µM of 4-(2-pyridylazo)resorcinol (PAR) was prepared in 50 mM Tris-HCl, 100 mM NaCl, pH 8 and stored protected from light in a 50 mL conical at 4 ^o^C.^2-3^ Concentrations of anti-TRAP and DnaJ were determined using BCA assay. A standard curve was prepared by incubating 0-20 µM ZnSO_4_·7H_2_O in 50 mM Tris, 100 mM NaCl, 10% glycerol, pH 7.4 with 5 M urea at 95 ^o^C for 30 min. Equivalent volumes of 200 µM PAR were then added right before absorbance measurement at 500 nm using BioTek Synergy 4 Multi-Detection Microplate Reader. Protein samples were also incubated in 5 M urea at 95 ^o^C for 30 min before mixing with equivalent volumes of 200 µM PAR. Protein samples were prepared so that the final concentrations when mixed with PAR were ~2 µM.

**Sequence similarity network (SSN) of NmaM**

The NmaM sequence was inputted into the Enzyme Function Initiative-Enzyme Similarity Tool (EFI-EST) to generate a sequence similarity network of 1000 sequences.^4-5^ An alignment score of 46 was initially used to create a 100% identity representative node network with 952 nodes. The network was visualized using the default Prefuse Force Directed layout in Cytoscape and further filtered to require >52% edge identity.

**Sequence analysis of cysimiditide precursors**

TceA (MAWGKKDTDDNNSKKDCGACDGDGGSWVYSDGQTQGKSHQRWVACQACGGSGKV) and TceM (WP_083948071.1) sequences from *Thermobifida cellulosilytica* were inputted into a BLASTP search in March 2024, and the results were manually examined for ORFs corresponding to cysimidtide precursors. Identified cysimiditides were combined with a list of cysimiditides previously identified through genome mining of imiditides^6^ to give a total list of 56 precursors (**Table S3**). A list of these precursor sequences starting from the first CXXC motif was aligned using Clustal Omega^7^ and then inputted into WebLogo^8^ to create a sequence logo. Precursors from *Salinactinospora qingdaonensis* (methyltransferase accession number GAA3754531.1) and *Streptomyces sp. SBT349* (methyltransferase accession number WP_049573986.1) were excluded from this WebLogo input as manual examination of these two sequences showed rare differences.

**Analysis of neighboring genes**

The list of methyltransferase accession numbers was inputted into webFlaGs^9^ (<https://server.atkinson-lab.com/webflags>) to look for gene conservation around cysimiditide BGCs, up to four flanking genes on each side of the methyltransferases. The 9 methyltransferase accession numbers from *Nocardiopsaceae bacterium* (MDA8370024.1), *Salinactinospora qingdaonensis* (GAA3731431.1), *Microbispora sp.* (MBX6383762.1), *Actinomadura rugatobispora* (BFE30814.1), *Salinactinospora qingdaonensis* (GAA3754531.1), *Candidatus Frankia californiensis* (SBW17366.1), *Frankia sp. CcI156* (ONH22856.1), *Frankia sp. CgIM4* (OFB40277.1), and *Frankia casuarinae* (ABD12601.1) were excluded, as they were not from the NCBI Reference Sequence database.

**Table S1.** Primers used in this study.

| **Primer Name** | **Primer Sequence (5’ -> 3’)** |
| --- | --- |
| OAZ1 | GCATGAGCTCATGACCTGGGGAAAGAAAGACA |
| OAZ2 | GCATAAGCTTGACTACGTGCTGCCATGAG |
| OAZ3 | GCATGGTCTCACATGCAACGGAACAGGAGAAC |
| OAZ4 | GCATGAAGCTTTTAGACTACGTGCTGCCATGAG |
| OAZ6 | GCATAGGATCCACCTGGGGAAAGAAAGACAG |
| OAZ7 | GCATGAAGCTTTCACCGCTGTTCTCCTGTT |
| OAZ9 | AATCTGTTCTCTGTGAGCCTCAA |
| OAZ11 | GCATGAATTCATTAAAGAGGAGAAATTAACTATGAGCGGATCG |
| OAZ17 | GCATTGCTAGCTTGGATTCTCACCAATAA |
| OAZ21 | GCATGGGATCCGCCTGGGGAAAGAAAGACACC |
| OAZ22 | GCATGAAGCTTTTACCCGTGCCGCCAGGA |
| OAZ23 | GCATGGTCTCACATGACATTCGAAGCAGCACTGCG |
| OAZ24 | GCATGAAGCTTTTACCCGTGCCG |
| OAZ25 | GCATAGGATCCACATTCGAAGCAGCACTGCG |
| OAZ27 | GCATAGGATCCGCCTGGGGAAAG |
| OAZ28 | GCATGAAGCTTTCAGACCTTCCCGCTGCC |
| OAZ29 | AGGCTCACAGAGAACAGATTGGTTCCGCCTGGGGAAAG |
| OAZ30 | CGCCGCACGCCTGACAT |
| OAZ31 | ATGTCAGGCGTGCGGC |
| OAZ41 | GCATAGGATCCGTGAGCGTCACCGTTCGGAA |
| OAZ42 | GCATGAAGCTTCTACACGGTCGTGTGCGGTC |
| OAZ43 | GCATAGGATCCATGGGTGACTGTTCGGGATGT |
| OAZ44 | GCATCGTCTCACATGAGGGACGTGAACAGGAC |
| OAZ45 | GCATCGTCTCAAGCTTCTACACGGTCGTG |
| OAZ50 | GCATGAAGCTTTCACGCCTTCCCCGTG |
| OAZ59 | GCGTCTGTCCGTTCGAGTAGAC |
| OAZ60 | GTCTACTCGAACGGACAGACGC |
| OAZ61 | GCATGGTCTCCCATGACCGAAACTCTCGTCTCCAC |
| OAZ62 | GGGGATGAACGGGTGGCG |
| OAZ63 | CGCCACCCGTTCATCCCC |
| OAZ64 | GTCGGCGGCCCGGTCCACGGGTCGGCCGCGGTGCCAGATCCGGTCGGGGA |
| OAZ65 | CACCGCGGCCGACCCGTGGACCGGGCCGCCGACCCGGACCGGTGGT |
| OAZ66 | GGCCTCGATCGGGCGGATCTGGTCGGCGGTCGCTTCTAGATCGTGCGG |
| OAZ67 | ACCGCCGACCAGATCCGCCCGATCGAGGCCGGGCAGGAGCTGCTCG |
| OAZ68 | GCATAGGATCCAGCGTCACCGTTCGGAAG |
| OAZ76 | GCATAGGATCCGGCACCACCACGAGCTTCAA |
| OAZ77 | GCATGAAGCTTCTAGGTGCTCTTCACGTTCCCGG |
| OAZ78 | CCCGACCCATTGCCACC |
| OAZ79 | GGTGGCAATGGGTCGGG |
| OAZ80 | TCATCTCCTTGTTCGCCCAAGG |
| OAZ81 | CCTTGGGCGAACAAGGAGATGA |
| OAZ82 | GCATAGGATCCACATAGGTTTTCTGTACTTAGAGTGAACG |
| OAZ83 | GCATGAAGCTTGCCCATTTTATCCAAATCGATTTAGACGC |
| OAZ84 | GCATAGGATCCGTCATTGCAACTGATGAT |
| OAZ85 | GACTGTAAGCTTTTACTTATTCAAATGCTTTTG |

**Table S2.** Plasmids used in this study.

| **Plasmid** | **Plasmid description** | **Vector** | **Cloning method** |
| --- | --- | --- | --- |
| pAZ1 | A region surrounding the *tfuAM* BGC (for cloning only) | pQE-80L | PCR on *T. fusca* genomic DNA using OAZ1 and OAZ2. Digested vector and insert with *Sac*I and *Hin*dIII. Ligation. |
| pAZ2 | TfuM (for *in vivo* coexpression)  (Sequence corresponds with that of QOS59465.1) | pRSF-Duet (in MCS1) | PCR on pAZ1 using OAZ3 and OAZ4. Digested vector with *Nco*I and *Hin*dIII and digested insert with *Bsa*I and *Hin*dIII. Ligation. |
| pAZ4 | His_6_-TfuA (for cloning only) | pQE-80L | PCR on pAZ1 using OAZ6 and OAZ7. Digested vector and insert with *Bam*HI and *Hin*dIII. Ligation. |
| pAZ12 | His_6_-TfuA | pQE-80L (modified His_6_-tag to start with MSGS) | PCR on pAZ4 using OAZ11 and OAZ7. Digested vector and insert *with Eco*RI and *Hin*dIII. Ligation. |
| pAZ18 | His_6_-TceA TceM (for cloning only) | pAZ12 | PCR on *T. cellulosilytica* genomic DNA using OAZ21 and OAZ22. Digested vector and insert with *Bam*HI and *Hin*dIII. |
| pAZ19 | TceM (for *in vivo* coexpression) | pRSF-Duet (in MCS1) | PCR on pAZ24 using OAZ23 and OAZ24. Digested vector with *Nco*I and *Hin*dIII and digested insert with *Bsa*I and *Hin*dIII. Ligation. |
| pAZ20 | His_6_-TceM (for *in vitro* reactions) | pQE-80L | PCR on pAZ24 using OAZ25 and OAZ24. Digested vector and insert with *Bam*HI and *Hin*dIII. Ligation. |
| pAZ22 | His_6_-TceA(G59C) | pAZ12 | PCR on pAZ24 using OAZ27 and OAZ28. Digested vector and insert with *Bam*HI and *Hin*dIII. Ligation. |
| pAZ23 | His_6_-SUMO-TceA(G156C) | pAZ12 | PCR on pLC53^6^ using OAZ11 and OAZ9 for insert 1 (SUMO tag) and PCR on pAZ24 using OAZ29 and OAZ28 for insert 2. Overlap PCR. Digested vector and final insert with *Eco*RI and *Hin*dIII. Ligation. |
| pAZ24 | His_6_-TceA(G59C) TceM (for cloning only) | pAZ12 | PCR on pAZ18 using OAZ27 and OAZ30 for insert 1 and PCR on pAZ18 using OAZ31 and OAZ24 for insert 2. Overlap PCR. Digested vector and final insert with *Bam*HI and *Hin*dIII. Ligation. |
| pAZ31 | His_6_-NalA NalM  (After the His_6_-tag, NalA starts as VSVTVRKAGGGSEGGERMGD before the first CXXC) | pAZ12 | PCR on *N. alba* genomic DNA using OAZ41 and OAZ42. Digested vector and insert with *Bam*HI and *Hin*dIII. Ligation. |
| pAZ32 | His_6_-NalA NalM  (After the His_6_-tag, NalA starts only with MGD before the first CXXC) | pAZ12 | PCR on pAZ31 using OAZ43 and OAZ42. Digested vector and insert with *Bam*HI and *Hin*dIII. Ligation. |
| pAZ33 | NalM (for *in vivo* coexpression)  (The initial MRD was included before the sequence of WP_234305832.1) | pRSF-Duet (in MCS1) | PCR on pAZ32 using OAZ44 and OAZ45. Digested vector with *Nco*I and *Hin*dIII and digested insert with *Bsm*BI. Ligation. |
| pAZ36 | His_6_-NalA  (After the His_6_-tag, NalA starts as SVTVRKAGGGSEGGERMGD before the first CXXC) | pAZ12 | PCR on pAZ31 using OAZ68 and OAZ50 (start codon GTG changed to ATG). Digested vector and insert with *Bam*HI and *Hin*dIII. Ligation. |
| pAZ44 | His_6_-TceA(D42N, G59C) | pAZ12 | PCR on pAZ22 using OAZ27 and OAZ59 for insert 1 and PCR on pAZ22 using OAZ60 and OAZ28 for insert 2. Overlap PCR. Digested vector and final insert with *Bam*HI and *Hind*III. Ligation. |
| pAZ45 | TfuM, deletion of residues 2-9 after the starting methionine  (for *in vivo* coexpression) | pRSF-Duet (in MCS1) | PCR on pAZ2 using OAZ61 and OAZ4. Digested vector with *Nco*I and *Hin*dIII and digested insert with *Bsa*I and *Hin*dIII. Ligation. |
| pAZ46 | TfuM, first 25 residues replaced with first 27 residues of TceM  (for *in vivo* coexpression) | pRSF-Duet (in MCS1) | PCR on pAZ19 using OAZ23 and OAZ62 for insert 1 and PCR on pAZ2 using OAZ63 and OAZ4 for insert 2. Overlap PCR. Digested vector with *Nco*I and *Hin*dIII and digested final insert with *Bsa*I and *Hin*dIII. Ligation. |
| pAZ47 | TfuM, residues 33-43 replaced with residues 35-45 of TceM  (for *in vivo* coexpression) | pRSF-Duet (in MCS1) | PCR on pAZ2 using OAZ3 and OAZ64 for insert 1 and PCR on pAZ2 using OAZ65 and OAZ4 for insert 2. Overlap PCR. Digested vector with *Nco*I and *Hin*dIII and digested final insert with *Bsa*I and *Hin*dIII. Ligation. |
| pAZ49 | TfuM, residues 242-251 replaced with residues 244-253 of TceM (for *in vivo* coexpression) | pRSF-Duet (in MCS1) | PCR on pAZ2 using OAZ3 and OAZ66 for insert 1 and PCR on pAZ2 using OAZ67 and OAZ4 for insert 2. Overlap PCR. Digested vector with *Nco*I and *Hin*dIII and digested final insert with *Bsa*I and *Hin*dIII. Ligation. |
| pAZ50a | A region surrounding anti-*trp* RNA-binding attentuation protein WP_003234807.1 (for cloning only) | pQE-80L | PCR on *Bacillus subtilis* strain 168 genomic DNA using OAZ82 and OAZ83. Digested vector and insert with *Bam*HI and *Hin*dIII. Ligation. |
| pAZ50 | His_6_-anti-*trp* RNA-binding attenuation protein (His_6_-anti-TRAP, WP_003234807.1) | pQE-80L | PCR on pAZ50a using OAZ84 and OAZ85. Digested vector and insert with *Bam*HI and *Hin*dIII. Ligation. |
| pAZ51 | His_6_-DnaJ(residues 146-244 of WP_068753762.1) | pQE-80L | PCR on *T. cellulosilytica* genomic DNA using OAZ76 and OAZ77. Digested vector and insert with *Bam*HI and *Hin*dIII. Ligation. |
| pAZ52 | His_6_-NalA(D46N)  (After the His_6_-tag, NalA starts as SVTVRKAGGGSEGGERMGD before the first CXXC) | pAZ12 | PCR on pAZ36 using OAZ11 and OAZ78 for insert 1 and PCR on pAZ36 using OAZ79 and OAZ17 for insert 2. Overlap PCR. Digested vector and insert with *Eco*RI and *Nhe*I. Ligation. |
| pAZ53 | His_6_-NalA(D53N)  (After the His_6_-tag, NalA starts as SVTVRKAGGGSEGGERMGD before the first CXXC) | pAZ12 | PCR on pAZ36 using OAZ11 and OAZ80 for insert 1 and PCR on pAZ36 using OAZ81 and OAZ17 for insert 2. Overlap PCR. Digested vector and insert with *Eco*RI and *Nhe*I. Ligation. |
| pAZ54 | His_6_-TceA | pAZ12 | PCR on pAZ18 using OAZ27 and OAZ28. Digested vector and insert with *Bam*HI and *Hin*dIII. Ligation. |
| pAZ55 | His_6_-SUMO-TceA | pAZ12 | PCR on pAZ23 using OAZ11 and OAZ9 for insert 1 and PCR on pAZ18 using OAZ29 and OAZ28 for insert 2. Overlap PCR. Digested vector and insert with *Eco*RI and *Hin*dIII. Ligation. |

### Supplementary Figures

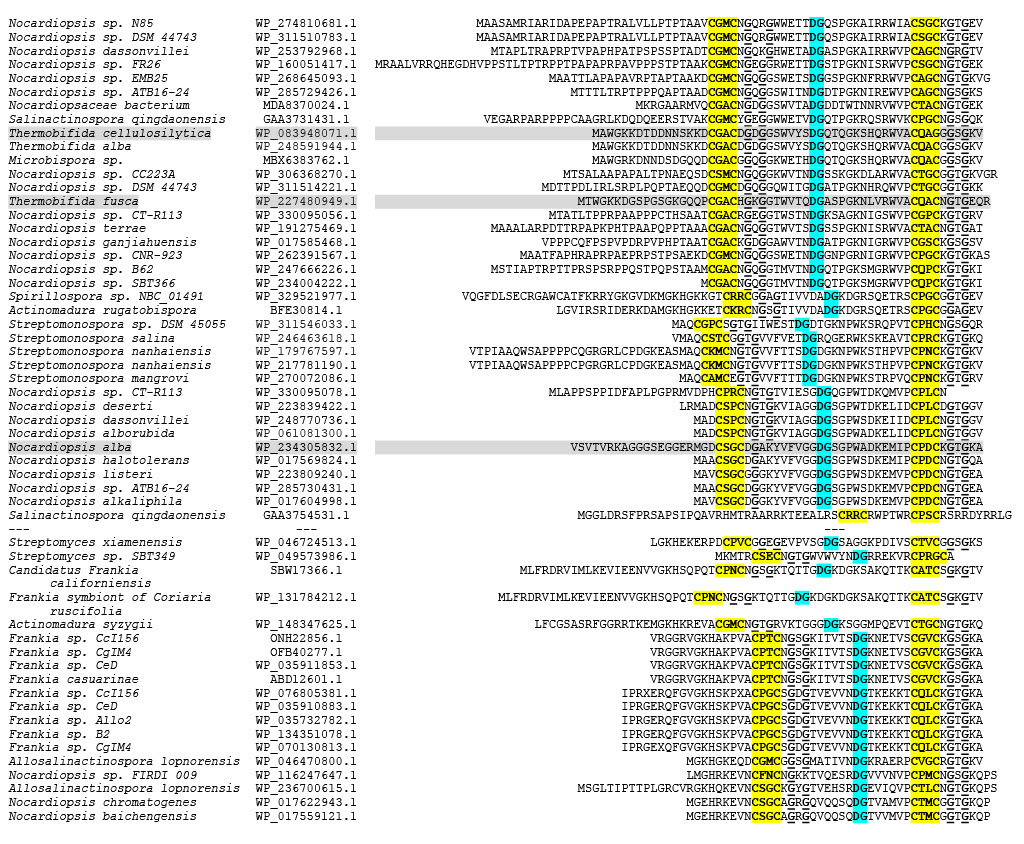

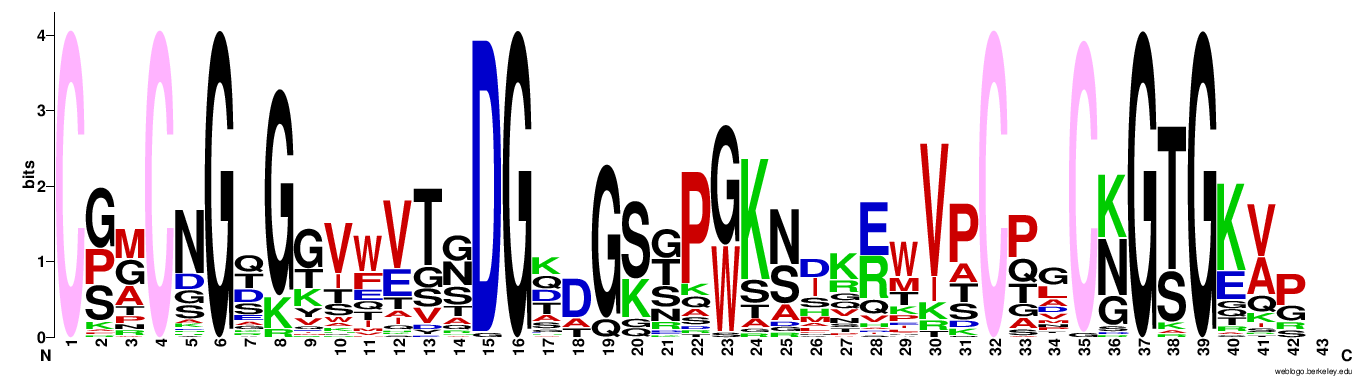

**Figure S1: Cysimiditides share conserved CXXCXGXG motifs with a potential aspartimidylation site between them.** **(top)** Cysimiditide precursor sequences have been aligned by the second CXXCXGXG motif (highlighted in yellow, with the Gs underlined). The potential aspartimidylation sites (DG, highlighted in blue) were identified via Clustal Omega alignment.^7^ The accession numbers listed correspond to the associated methyltransferases. Cysimiditides in the top half were identified via BLASTP of the precursor and methyltransferase sequences from *Thermobifida cellulosilytica*; those in the bottom half were identified previously through imiditide genome mining.^6^ **(bottom)** Sequence logo^8^ of cysimiditides showing conserved residues, namely, the cysteine motifs and the putative aspartimidylation site.

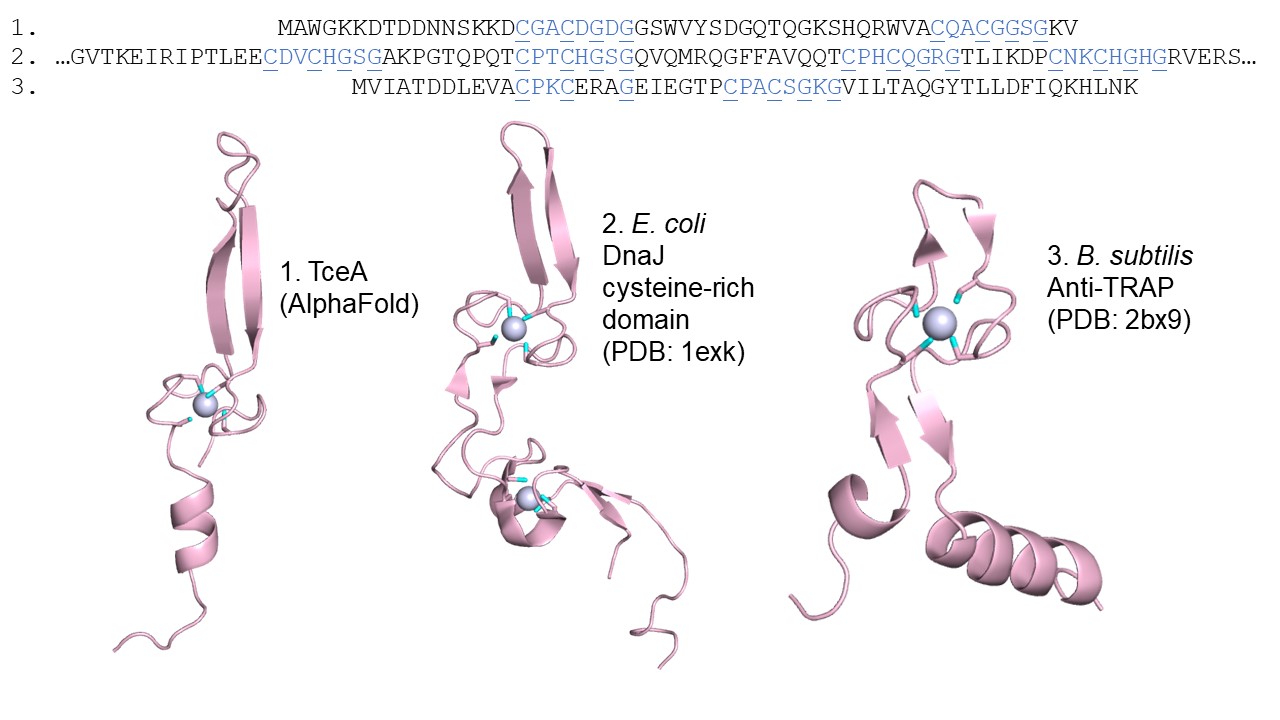

**Figure S2: Proteins with CXXCXGXG zinc-binding motifs.** Sequences and crystal structures of the cysteine-rich region of *E. coli* DnaJ (PDB: 1exk)^10^ and *Bacillus subtilis* anti-*trp* RNA-binding attenuation protein (anti-TRAP) (PDB: 2bx9)^11^ show bound zinc ions, with each ion coordinated by four cysteines. The structure of TceA was obtained by inputting the TceA precursor sequence into the AlphaFold 3 server (with the second cysteine motif corrected from CQAG to CQAC) and modeling it with a Zn^2+^ ion.^12^ Analogous to the DnaJ and anti-TRAP structures, the AlphaFold 3 model of TceA has the zinc ion coordinated by four cysteines.

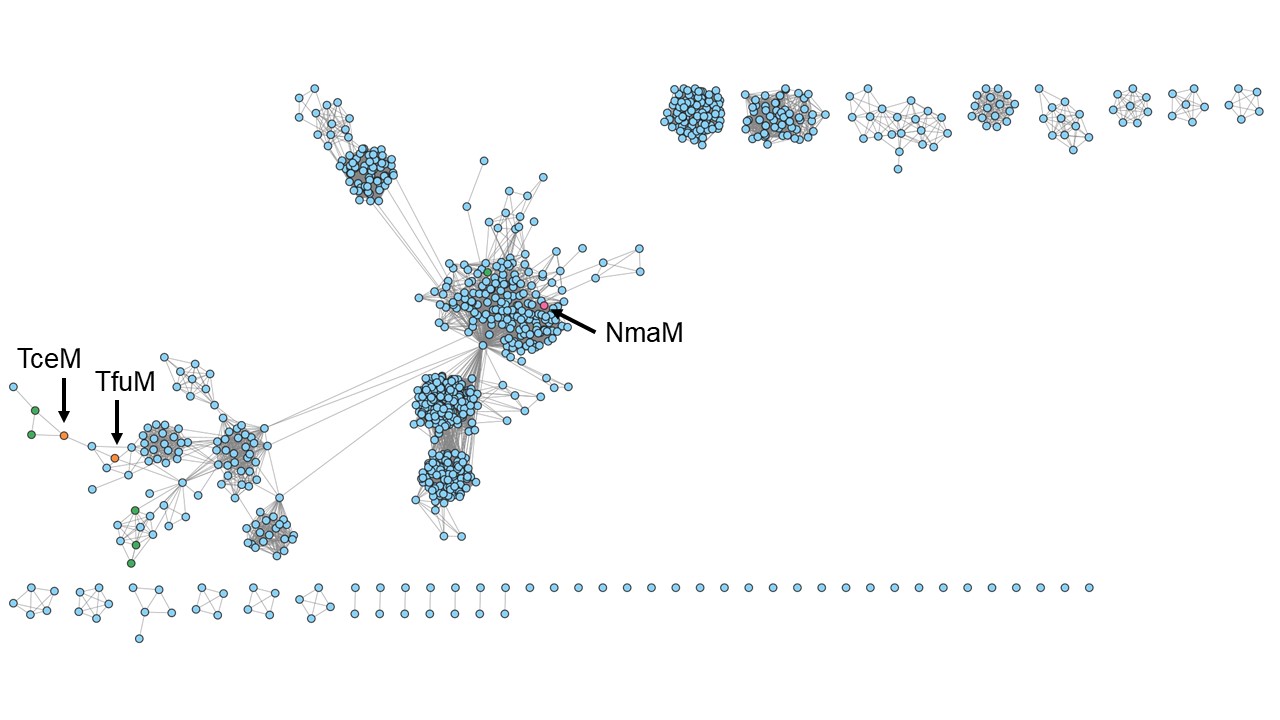

**Figure S3: Sequence similarity network (SSN) of the imiditide methyltransferase NmaM.** The SSN of 1000 sequences was initially created with an alignment score of 46 and further filtered in Cytoscape to require >52% edge identity.^4-5^ The nodes representing NmaM from *Nonomuraea maritima*, the cysimiditide methyltransferase TceM from *Thermobifida cellulosilytica*, and the cysimiditide methyltransferase TfuM from *Thermobifida fusca* are colored orange. Green nodes represent other cysimiditide methyltransferases from Table S3.

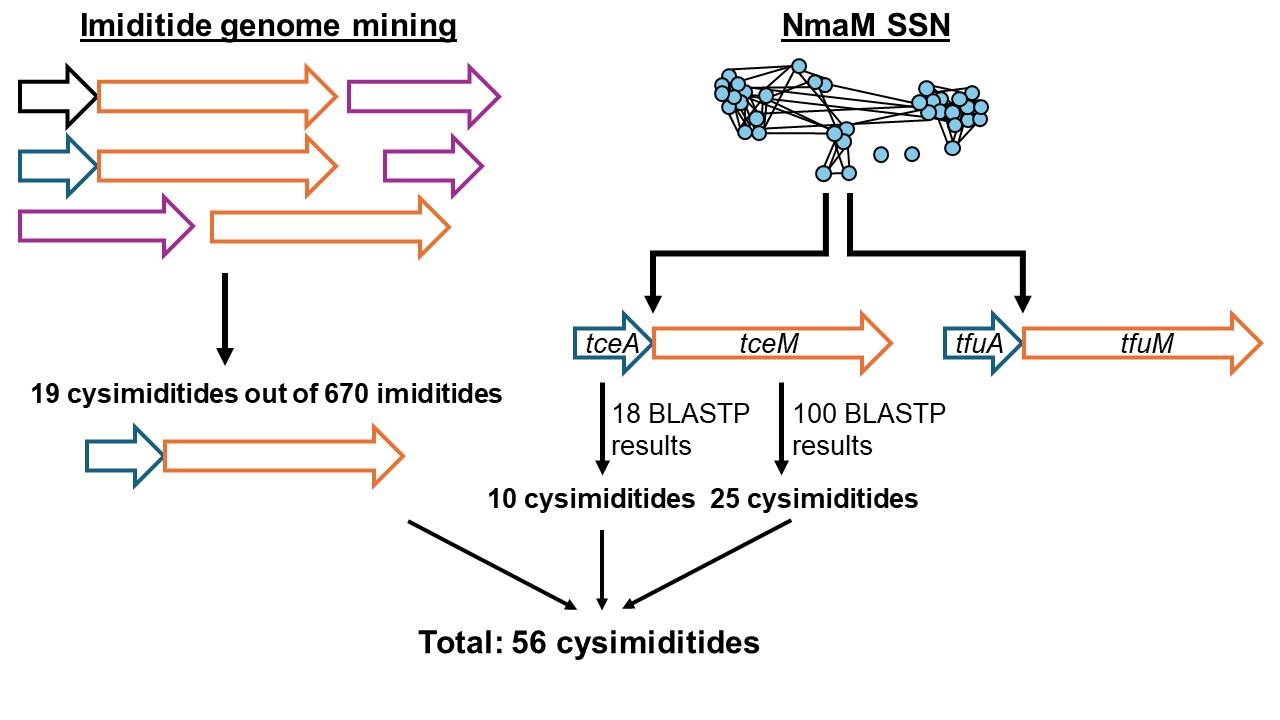

**Figure S4: Computational methods used to identify putative cysimiditide BGCs.** Manual examination of imiditide genome mining results^6^ revealed 19 cysimiditides. The sequence similarity network (SSN) of a characterized imiditide methyltransferase NmaM (Figure S3) contained two cysimiditides from *Thermobifida cellulosilytica* and *Thermobifida fusca*, organisms from which we have previously identified RiPPs.^13-15^ Using the *T. cellulosilytica* cysimiditide precursor TceA and its associated methyltransferase TceM as inputs for BLASTP led to discovery of more cysimiditides that were not identified through imiditide genome mining. In total, 56 cysimiditide BGCs including *tceAM* and *tfuAM* were found.

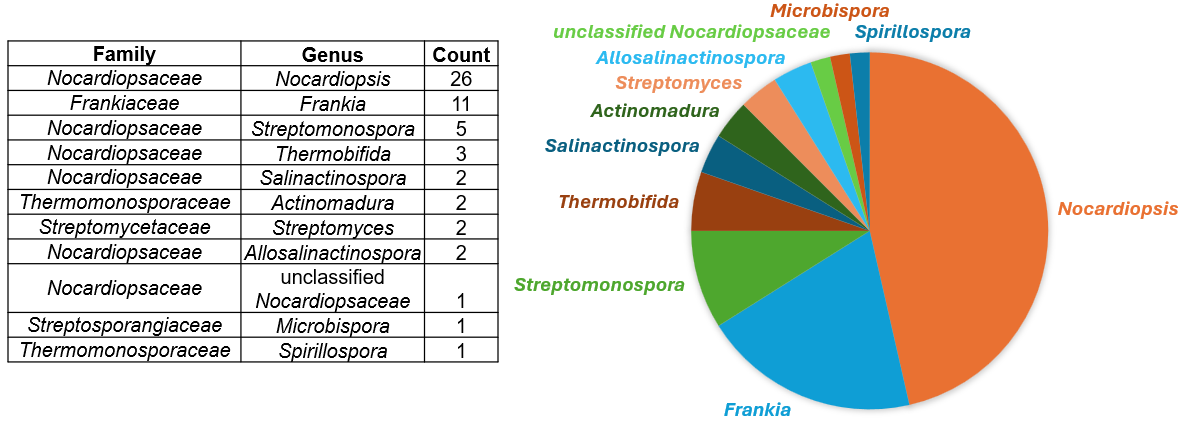

**Figure S5: Distribution of putative cysimiditide BGCs by bacteria family and genus.** The majority are from *Nocardiopsis* and *Frankia*.

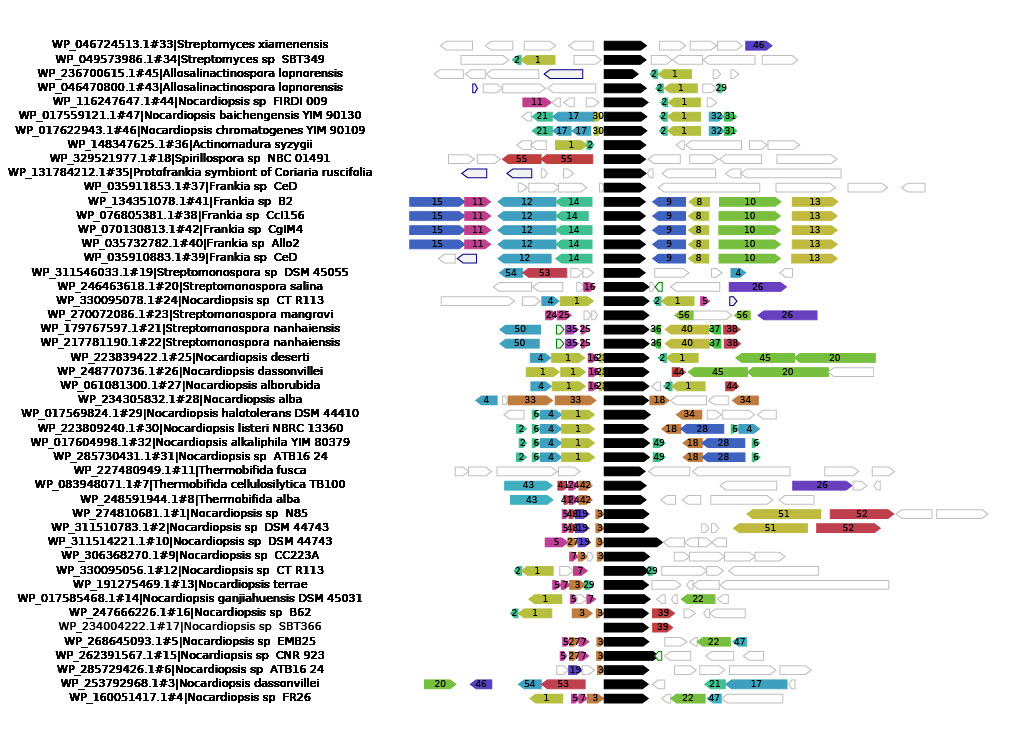

29. BldC family transcriptional regulator

31. Helix-turn-helix domain-containing protein

33. Methyltransferase domain-containing protein

34. TetR/AcrR family transcriptional regulator

37. Metalloregulator ArsR/SmtB family transcription factor

38. Arsenate reductase ArsC

39. ATP-binding protein

40. ACR3 family arsenite efflux transporter

45. Rhamnulokinase family protein

46. Pentapeptide repeat-containing protein

49. NIPSNAP family protein

50. D-alanyl-D-alanine carboxypeptidase family protein

51. serine hydrolase domain-containing protein

52. OmpA family protein

53. Helix-turn-helix transcriptional regulator

54. HD domain-containing protein

55. XRE family transcriptional regulator

56. MerR family transcriptional regulator

1. Helix-turn-helix transcriptional regulator

2. DUF397 domain-containing protein

3-7, 12, 14, 16, 19, 23-25, 27, 30, 32, 35-36, 41-44, 47-48. Hypothetical protein

8. Toxin-antitoxin system HicB family antitoxin

9. DUF4097 family beta strand repeat-containing protein

10. Serine/threonine protein kinase

11. Response regulator transcription factor

13. Glycerophosphodiester phosphodiesterase

15. HAMP domain-containing sensor histidine kinase

17. Asparagine synthase-related protein

18. HIT family protein

20. Bifunctional aldolase/short-chain dehydrogenase

21. GNAT family N-acetyltransferase

22. Peptidoglycan recognition family protein

26. Aldehyde dehydrogenase family protein

28. ThiF family adenylyltransferase

**Figure S6. Neighboring genes near putative cysimiditide BGCs.** webFlaGs^9^ was used to generate this figure and a list of the most frequently occuring genes near cysimiditide methyltransferases (shown as black arrows). Green-outlined arrows represent RNA coding genes, and blue-outlined arrows represent pseudogenes. Uncolored arrows represent genes that did not appear in the list of most frequently occuring genes. Helix-turn-helix transcriptional regulators (labeled “1” in figure), DUF397 domain-containing proteins (labeled “2” in figure), and hypothetical proteins are most frequently found near putative cysimiditide BGCs.

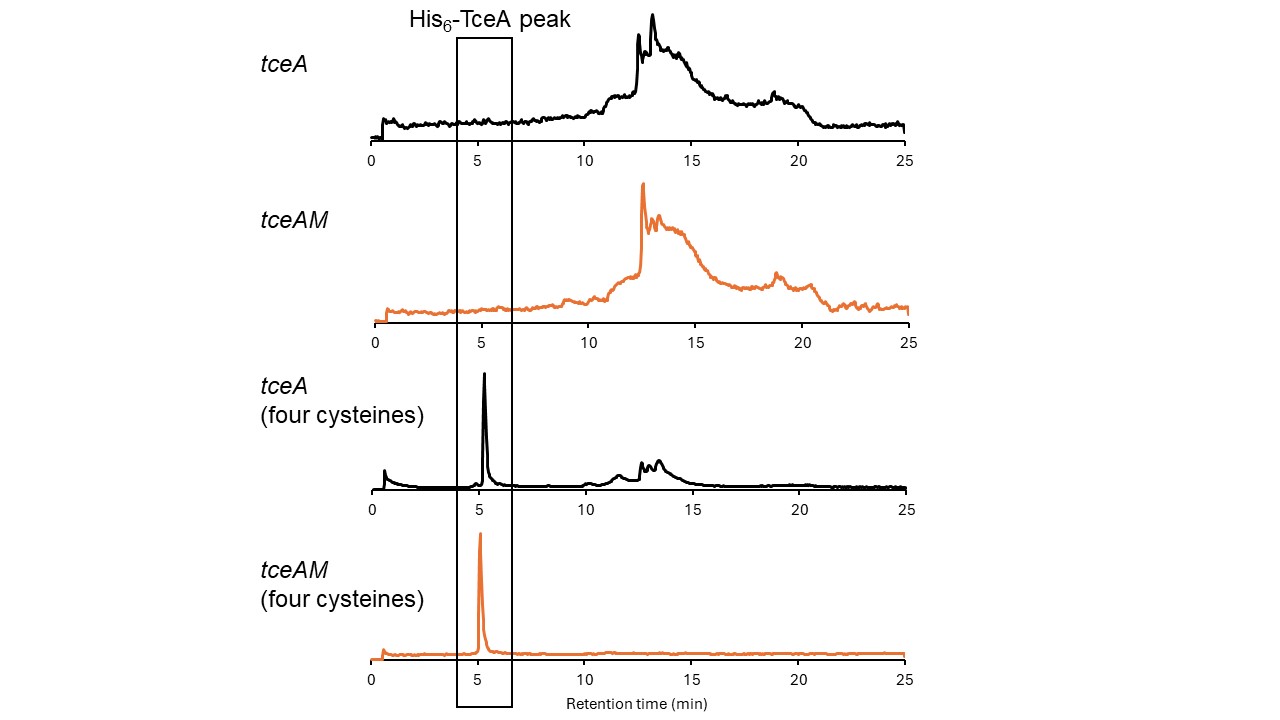

**Figure S7: His_6_-TceA cannot be expressed unless all four cysteines are present.** The top two total ion chromatograms are expressions of TceA with CQAG as the second cysteine motif, while the bottom two chromatograms are expressions of TceA with CQAC as the second cysteine motif. The total ion chromatograms did not contain an expected peak corresponding to the mass of His_6_-TceA at ~5 min retention time unless all four cysteines were present. This suggests that the cysteine motifs contribute to the stability of the protein *in vivo*.

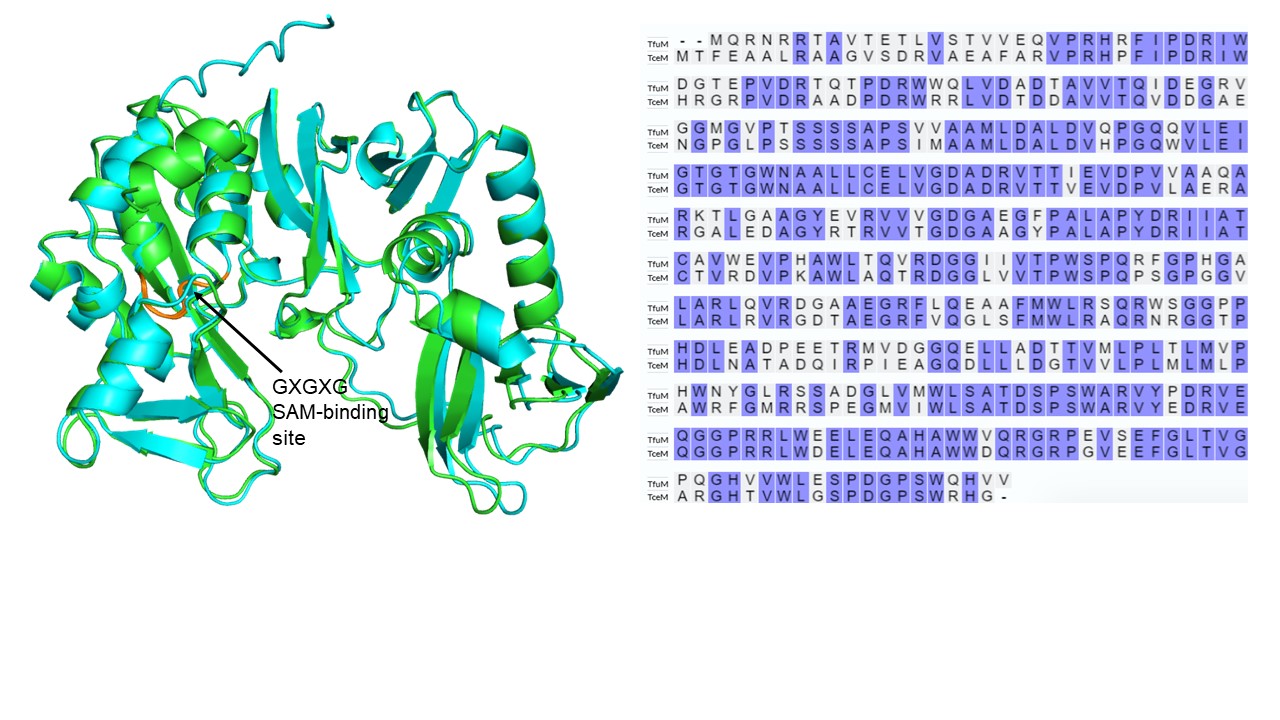

**Figure S8: AlphaFold structures and sequence alignment of TceM and TfuM.** Structures were predicted by the AlphaFold 3 server.^12^ TceM is shown in green, and TfuM is shown in cyan. The SAM-binding site (GXGXG) is shown in orange. The two methyltransferases share 68% sequence identity. Sequence alignment was done through inputting sequences into the UniProt Align tool.

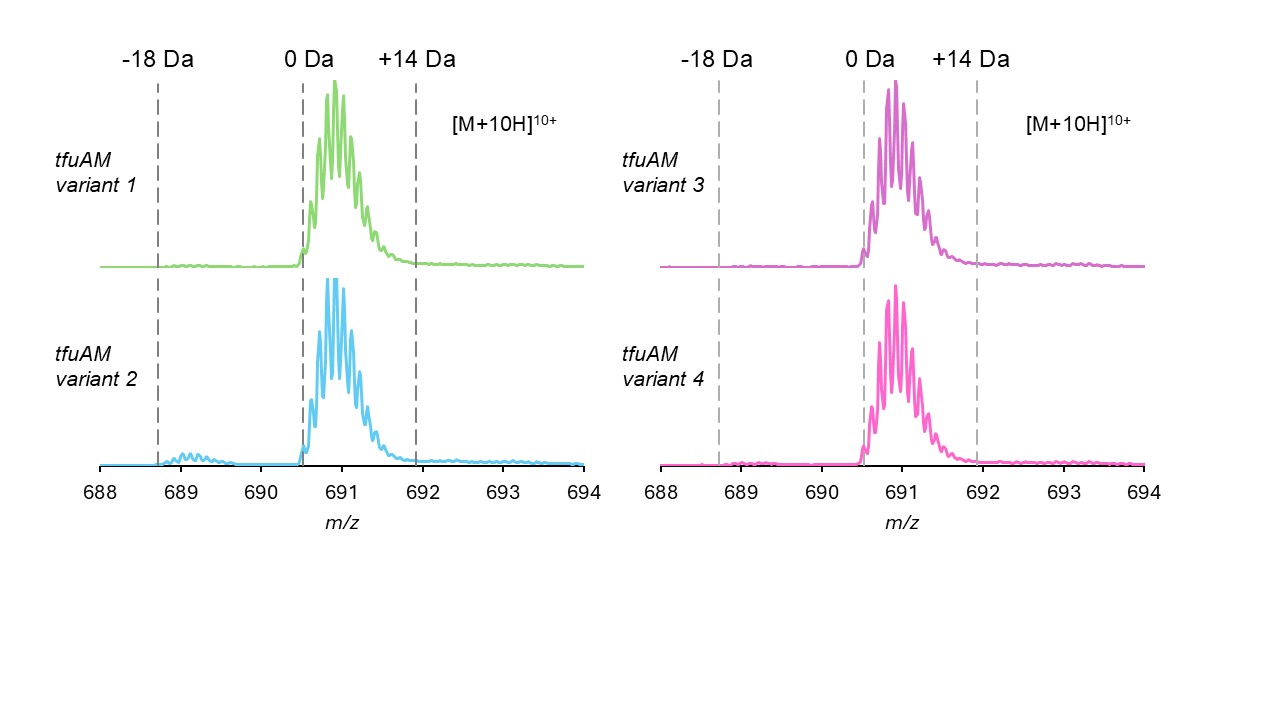

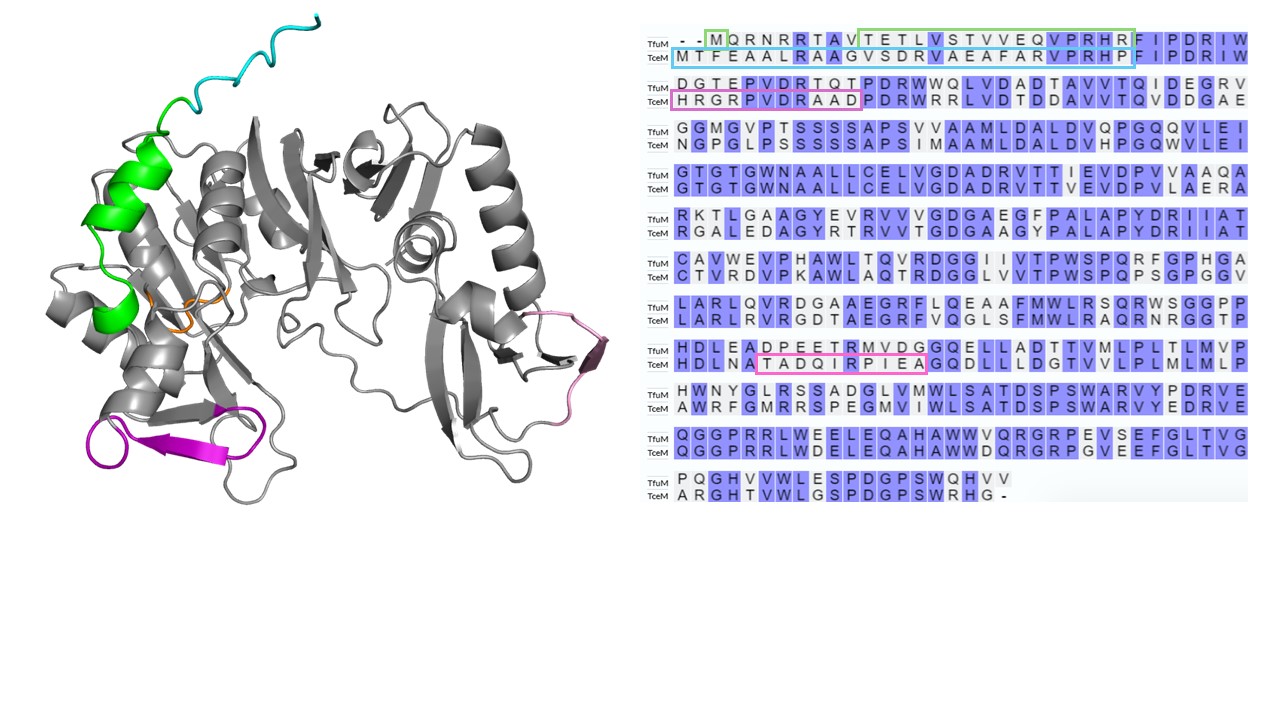

| **TfuM variant 1:** Deletion of residues 2-9 after the starting methionine | MTETLVSTVVEQVPRHRFIPDRIWDGTEPVDRTQTPDRWWQLVDADTAVVTQIDEGRVGGMGVPTSSSSAPSVVAAMLDALDVQPGQQVLEIGTGTGWNAALLCELVGDADRVTTIEVDPVVAAQARKTLGAAGYEVRVVVGDGAEGFPALAPYDRIIATCAVWEVPHAWLTQVRDGGIIVTPWSPQRFGPHGALARLQVRDGAAEGRFLQEAAFMWLRSQRWSGGPPHDLEADPEETRMVDGGQELLADTTVMLPLTLMVPHWNYGLRSSADGLVMWLSATDSPSWARVYPDRVEQGGPRRLWEELEQAHAWWVQRGRPEVSEFGLTVGPQGHVVWLESPDGPSWQHVV |
| --- | --- |
| **TfuM variant 2:**  First 25 residues replaced with first 27 residues of TceM | MTFEAALRAAGVSDRVAEAFARVPRHPFIPDRIWDGTEPVDRTQTPDRWWQLVDADTAVVTQIDEGRVGGMGVPTSSSSAPSVVAAMLDALDVQPGQQVLEIGTGTGWNAALLCELVGDADRVTTIEVDPVVAAQARKTLGAAGYEVRVVVGDGAEGFPALAPYDRIIATCAVWEVPHAWLTQVRDGGIIVTPWSPQRFGPHGALARLQVRDGAAEGRFLQEAAFMWLRSQRWSGGPPHDLEADPEETRMVDGGQELLADTTVMLPLTLMVPHWNYGLRSSADGLVMWLSATDSPSWARVYPDRVEQGGPRRLWEELEQAHAWWVQRGRPEVSEFGLTVGPQGHVVWLESPDGPSWQHVV |
| **TfuM variant 3:** Residues 33-43 replaced with residues 35-45 of TceM | MQRNRRTAVTETLVSTVVEQVPRHRFIPDRIWHRGRPVDRAADPDRWWQLVDADTAVVTQIDEGRVGGMGVPTSSSSAPSVVAAMLDALDVQPGQQVLEIGTGTGWNAALLCELVGDADRVTTIEVDPVVAAQARKTLGAAGYEVRVVVGDGAEGFPALAPYDRIIATCAVWEVPHAWLTQVRDGGIIVTPWSPQRFGPHGALARLQVRDGAAEGRFLQEAAFMWLRSQRWSGGPPHDLEADPEETRMVDGGQELLADTTVMLPLTLMVPHWNYGLRSSADGLVMWLSATDSPSWARVYPDRVEQGGPRRLWEELEQAHAWWVQRGRPEVSEFGLTVGPQGHVVWLESPDGPSWQHVV |
| **TfuM variant 4:** Residues 242-251 replaced with residues 244-253 of TceM | MQRNRRTAVTETLVSTVVEQVPRHRFIPDRIWDGTEPVDRTQTPDRWWQLVDADTAVVTQIDEGRVGGMGVPTSSSSAPSVVAAMLDALDVQPGQQVLEIGTGTGWNAALLCELVGDADRVTTIEVDPVVAAQARKTLGAAGYEVRVVVGDGAEGFPALAPYDRIIATCAVWEVPHAWLTQVRDGGIIVTPWSPQRFGPHGALARLQVRDGAAEGRFLQEAAFMWLRSQRWSGGPPHDLEATADQIRPIEAGQELLADTTVMLPLTLMVPHWNYGLRSSADGLVMWLSATDSPSWARVYPDRVEQGGPRRLWEELEQAHAWWVQRGRPEVSEFGLTVGPQGHVVWLESPDGPSWQHVV |

**Figure S9: Co-expressing TfuA with TfuM variants containing sequence substitutions from TceM did not result in aspartimidylation.** The AlphaFold 3 structure of TfuM is shown with the substituted sections colored according to the sequence alignment on the right.^12^ Amino acid sequences of each TfuM variant/chimera are shown below.

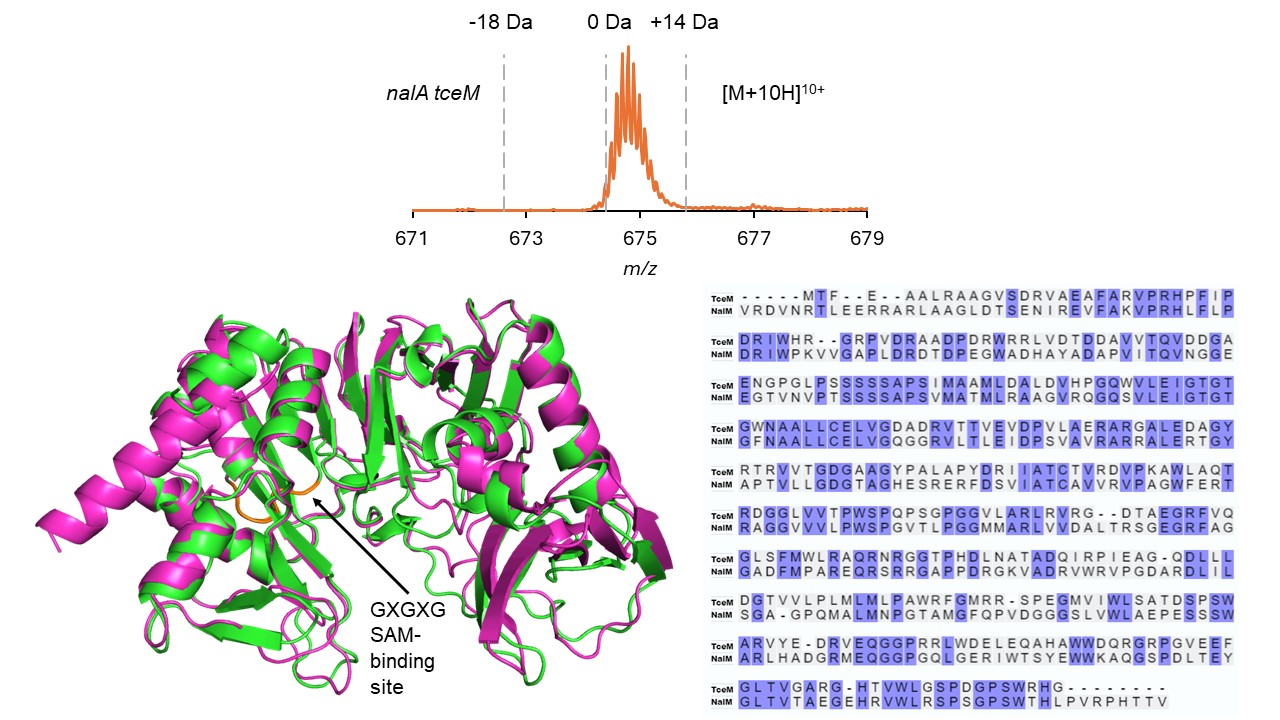

**Figure S10: TceM cannot modify His_6_-NalA *in vivo*.** The co-expression of His_6_-NalA with TceM did not show modification. TceM and NalM share 48% sequence identity but are structurally similar. Sequence alignment was done through inputting sequences into the UniProt Align tool. Structures were predicted by the AlphaFold 3 server.^12^ TceM is shown in green, and NalM is shown in pink. The SAM-binding site (GXGXG) is shown in orange.

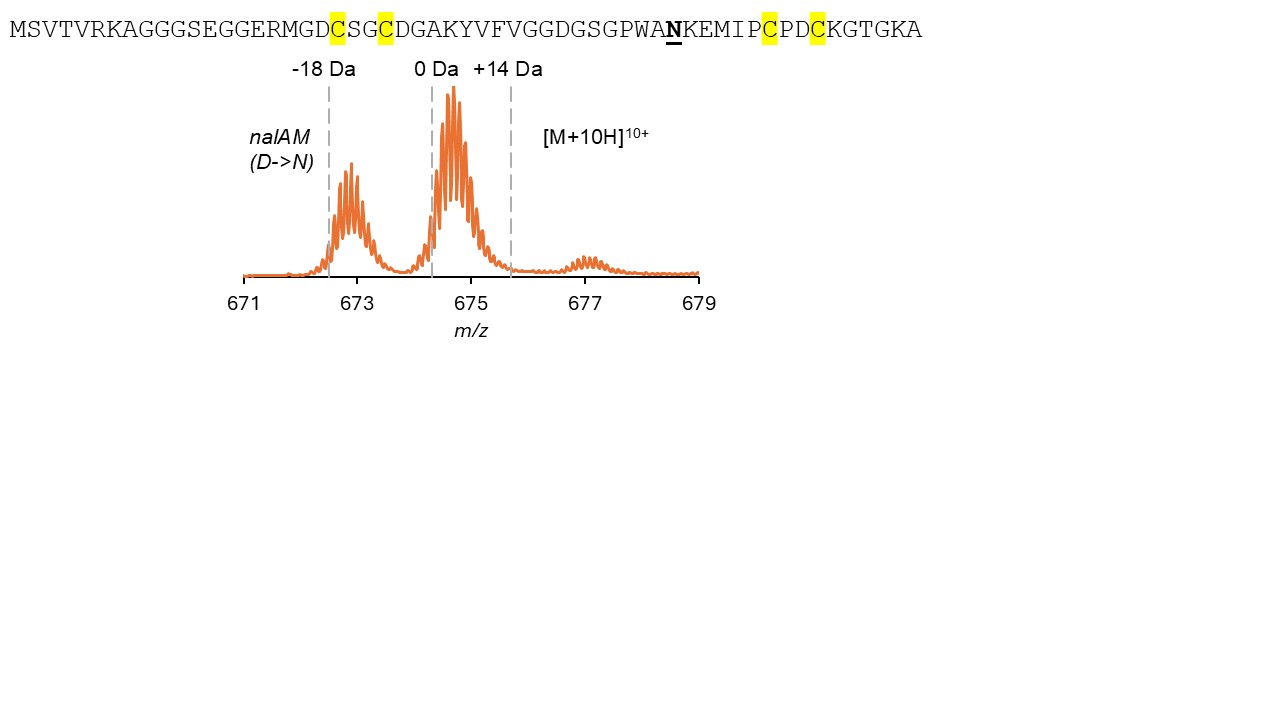

**Figure S11: Mutating the non-conserved Asp in His_6_-NalA to Asn did not prevent aspartimidylation from occurring.** Thus, this residue (shown bolded and underlined) is not the one being modified.

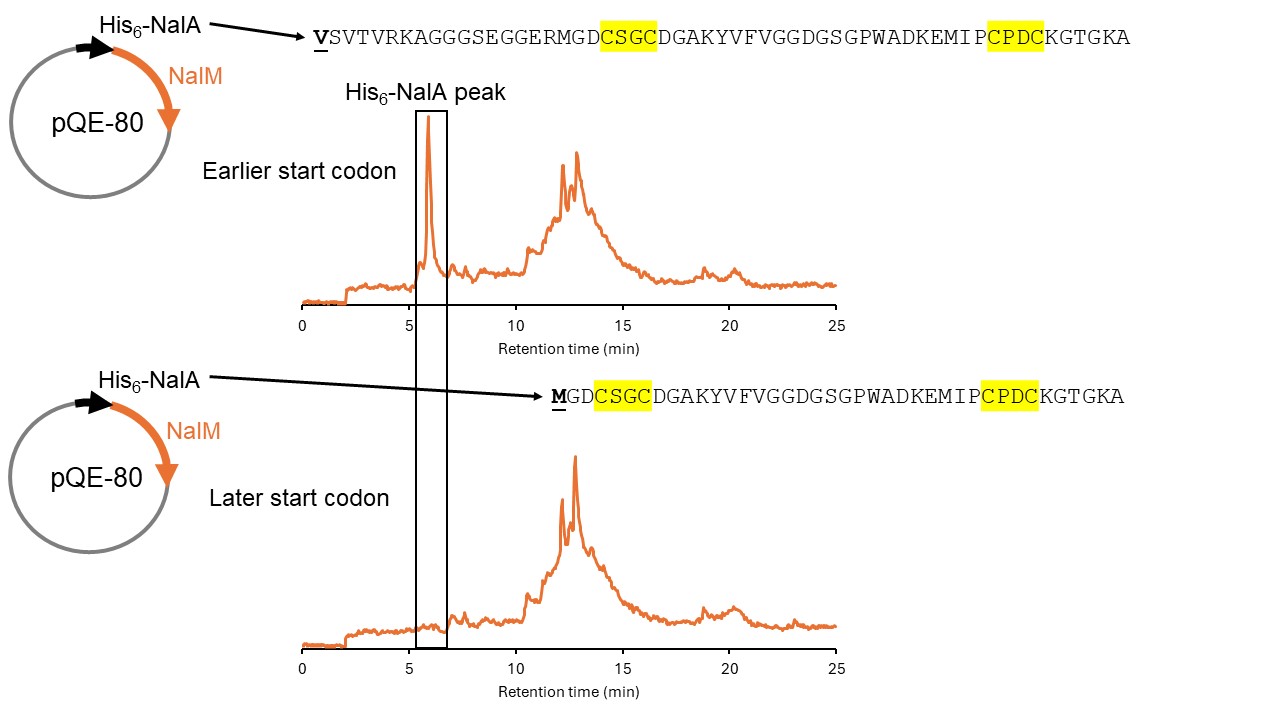

**Figure S12: His_6_-NalA was not expressed when the leader peptide was truncated to a later start codon.** The BGC consisting of *nalA* and *nalM* was placed in one plasmid under a T5 promoter with a N-terminal His_6_-tag. Two different start codons (shown underlined and bolded) were chosen for *nalA*. The total ion chromatogram for the later start codon beginning three amino acids before the first CXXC motif (bottom) did not contain an expected peak corresponding to the mass of His_6_-NalA at ~6 min retention time.

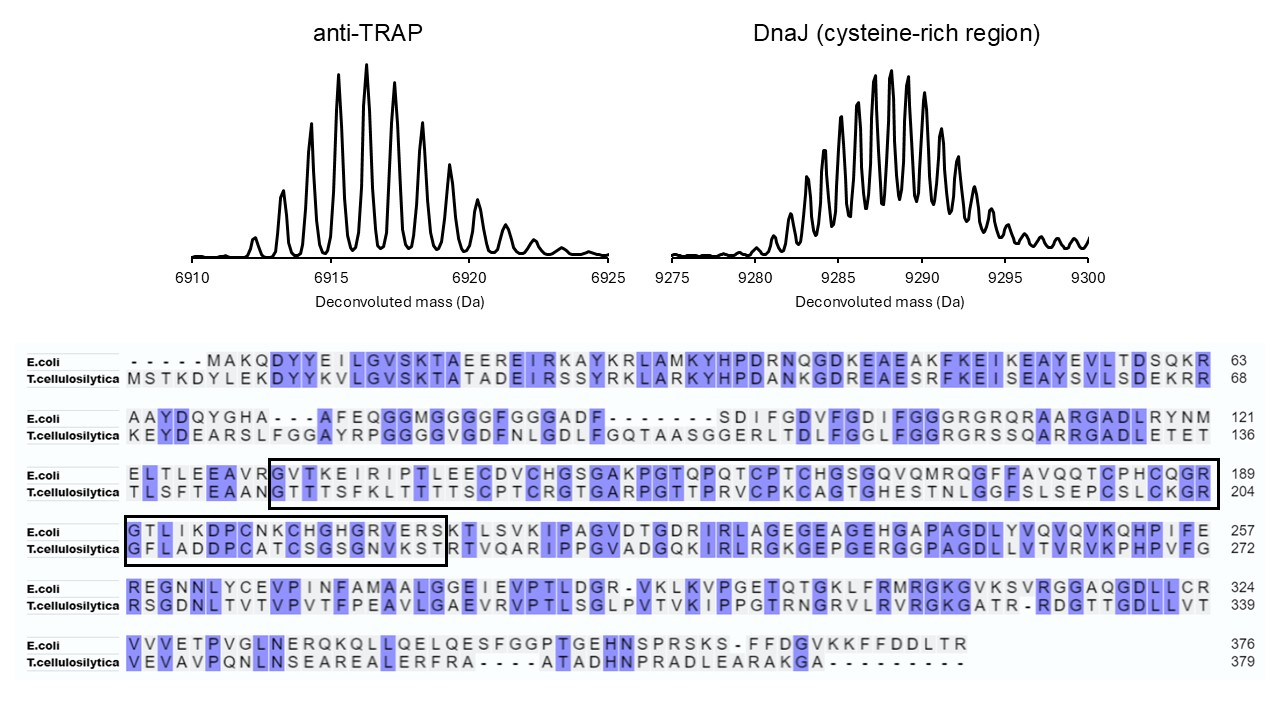

**Figure S13: Native purification of *Bacillus subtilis* anti-TRAP and the cysteine-rich region of *Thermobifida cellulosilytica* DnaJ.** The cysteine-rich region of *T. cellulosilytica* DnaJ was determined through sequence alignment with *E. coli* DnaJ, shown on the bottom outlined in black.^10^ Expected monoisotopic masses were 6912.37 Da for anti-TRAP and 9284.24 Da for DnaJ. The DnaJ purification also contained a species with two disulfide bonds present, which suggests the partial loss of one of the two zinc ions.

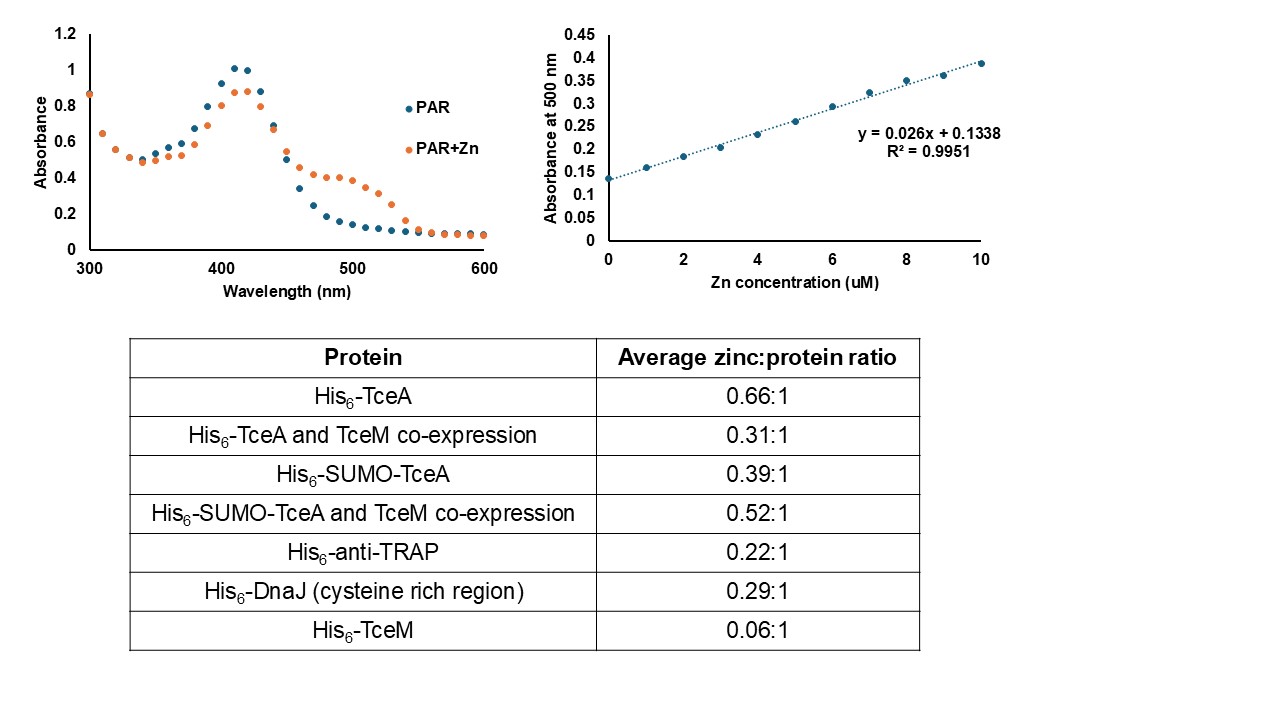

**Figure S14: PAR zinc-binding assay.** **(top)** The zinc-binding reagent 4-(2-pyridylazo)resorcinol (PAR) shows a maximal difference in absorbance at 500 nm when bound to zinc.^2^ A representative standard curve for absorbances at 500 nm is shown for 100 µM PAR mixed with 1-10 µM ZnSO_4_·7H_2_O. **(bottom)** TceA samples had greater zinc:protein ratios than the negative control TceM. Protein samples were incubated in 5 M urea at 95 ^o^C for 30 min before being mixed with PAR (final concentrations ~2 µM protein and 100 µM PAR in 100 uL). Anti-TRAP and DnaJ are known zinc-binding proteins, while TceM was not expected to bind zinc. The assay was repeated three times, and the average zinc:protein ratios are shown.

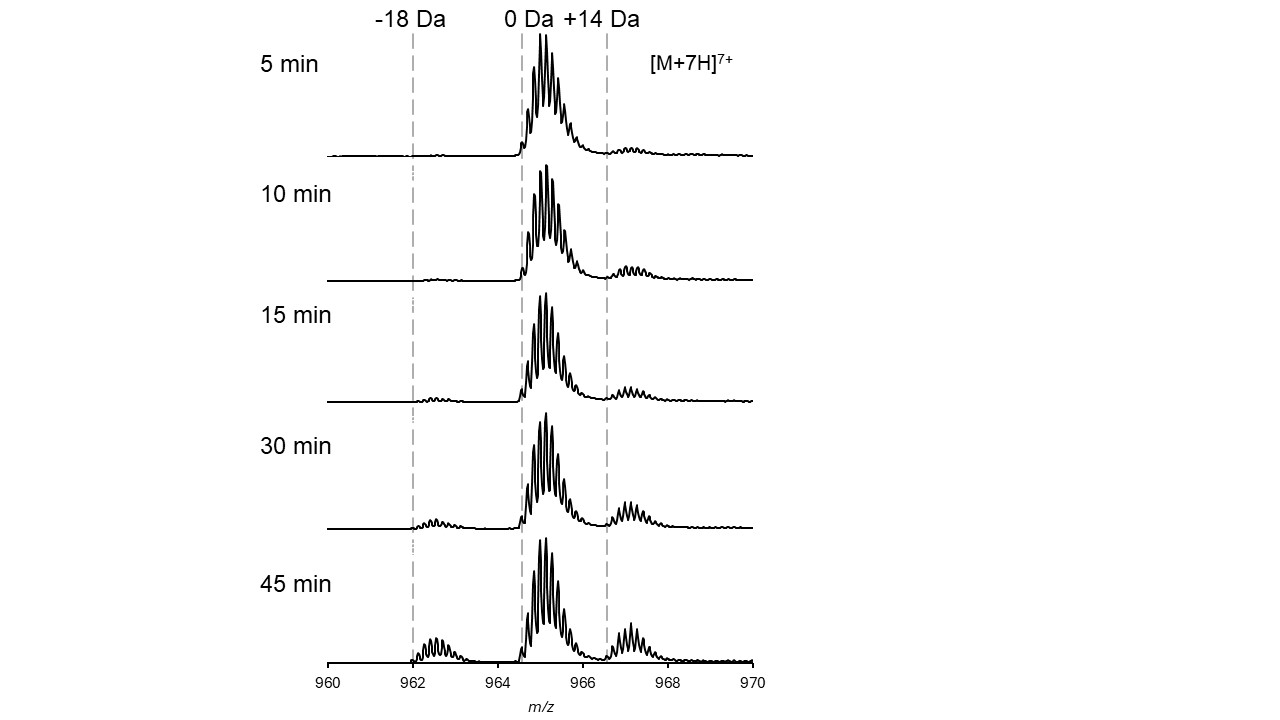

**Figure S15: *In vitro* reactions of His_6_-TceA with His_6_-TceM between 5 to 45 minutes.** Methylated species (+14 Da) appeared within 5 minutes, and aspartimidylated species (-18 Da) started to appear at 15 minutes. Reactions were performed with 5 µM His_6_-TceA and 1 µM His_6_-TceM in 50 mM Tris-HCl, pH 7.4, 400 µM SAM, 1 mM DTT, and 100 µM ZnCl_2_ at 37 ^o^C.

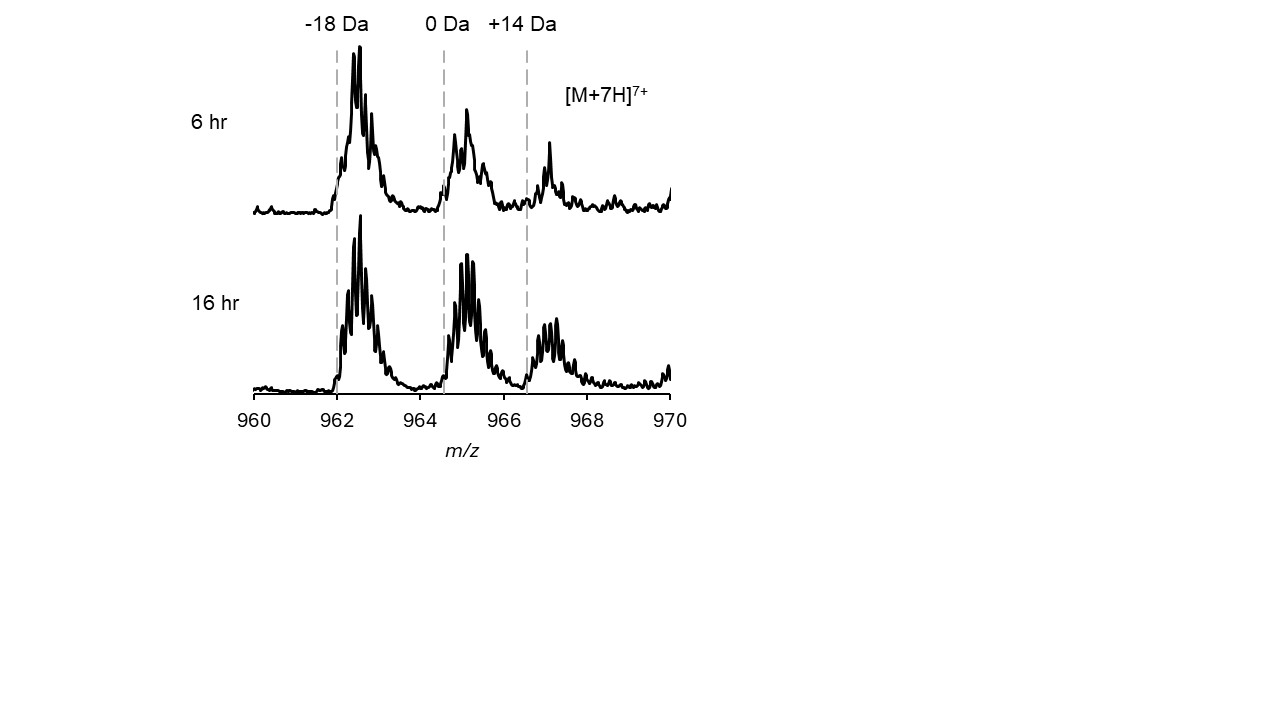

**Figure S16: *In vitro* reactions of His_6_-TceA with His_6_-TceM for 6 h and 16 h.** The amount of aspartimidylated species (-18 Da) did not substantially increase beyond that seen after 3 h of reaction (Figure 4A). Reactions were performed with 5 µM His_6_-TceA and 1 µM His_6_-TceM in 50 mM Tris-HCl, pH 7.4, 400 µM SAM, 1 mM DTT, and 100 µM ZnCl_2_ at 37 ^o^C.

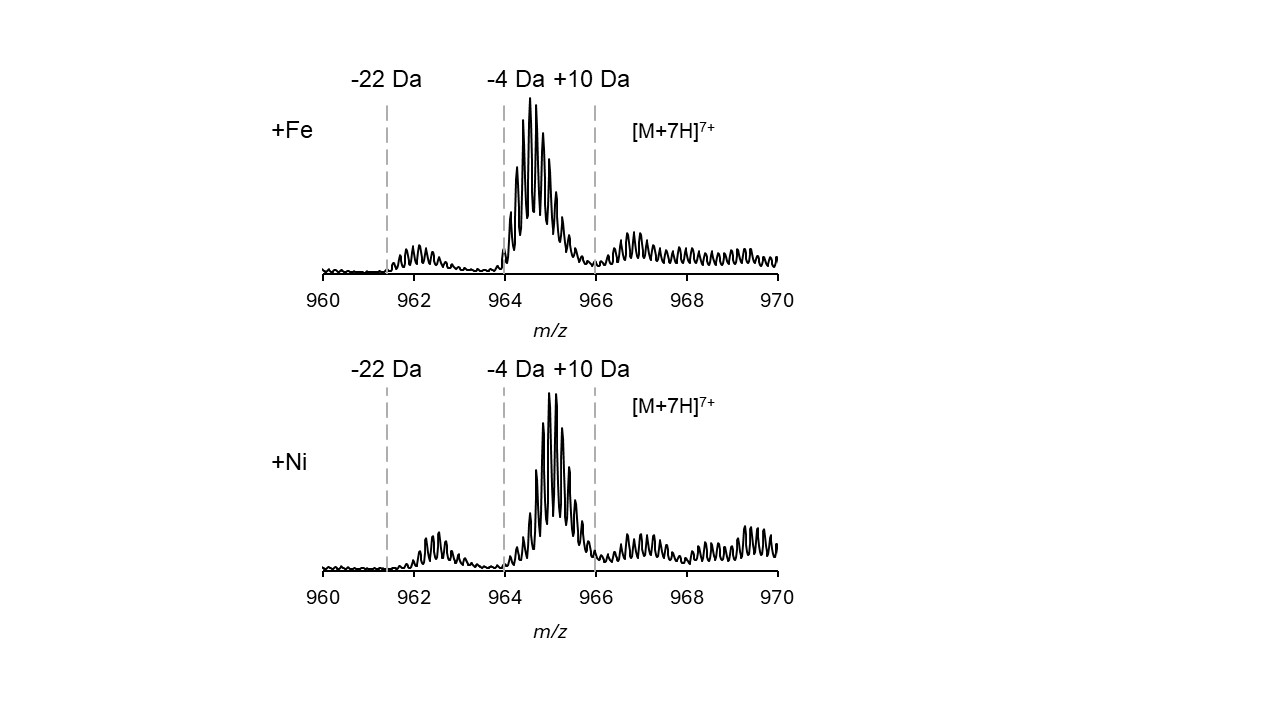

**Figure S17: Adding Fe^2+^ or Ni^2+^ to Zn^2+^-removed His_6_-TceA did not restore aspartimidylation by TceM.** *In vitro* reactions were performed with His_6_-TceA that was urea purified, incubated with 3 mM EDTA for 15 min at 37 ^o^C, and then desalted to remove both zinc and EDTA. 100 µM Fe(II) sulfate heptahydrate or Ni(II) sulfate hexahydrate was added to the reaction, and the reaction proceeded for 3 h at 37 ^o^C. Modification was not restored by adding these metals. Species with two disulfide bonds were observed. -22 Da corresponds to aspartimidylated species with two disulfide bonds, -4 Da corresponds to species with two disulfide bonds (either unmodified, or aspartimidylated and then hydrolyzed), and +10 Da corresponds to methylated species with two disulfide bonds.

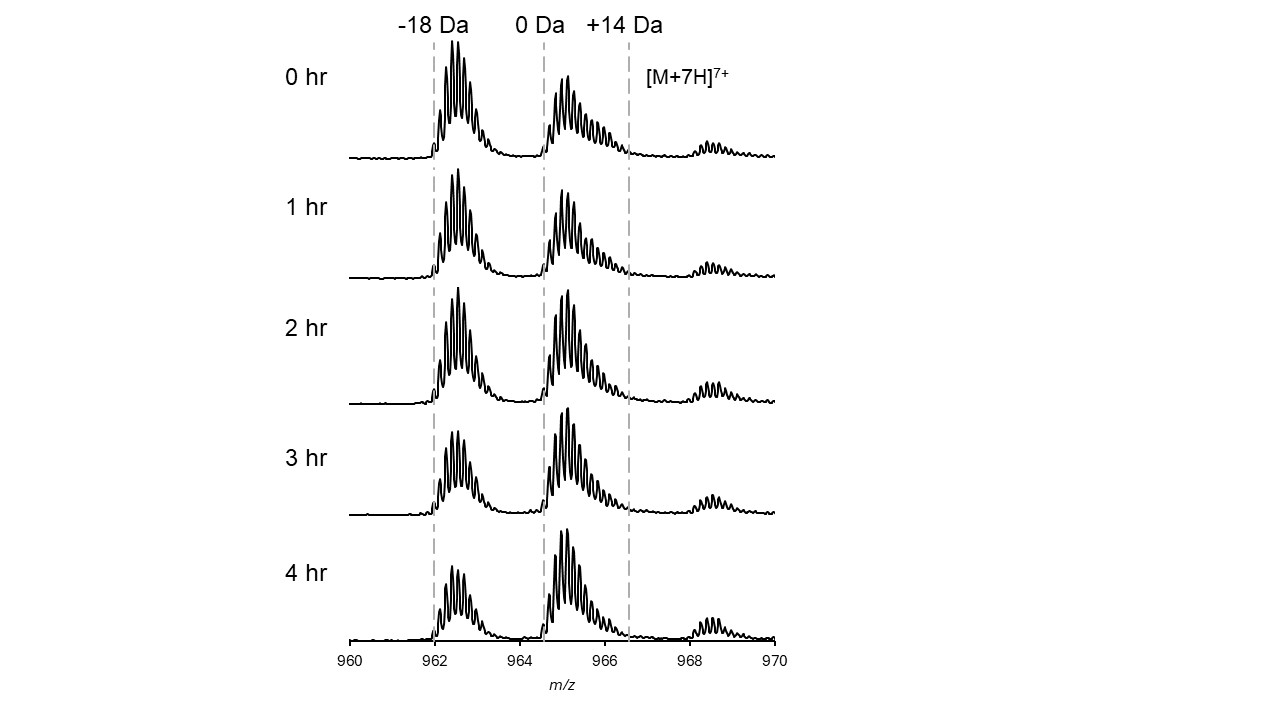

**Figure S18: Stability analysis of the natively purified His_6_-TceA and TceM co-expression product over 4 h at room temperature.** The aspartimidylated species (-18 Da) hydrolyzed over the course of 4 h in 50 mM Tris, 100 mM NaCl, 10% glycerol, pH 7.4. The 0 Da species represents either unmodified TceA or TceA that was aspartimidylated and then hydrolyzed, while +14 Da corresponds to methylated species.

### Supplementary Tables

**Table S3.** A list of the NCBI accession numbers corresponding to putative cysimiditide BGCs that were identified through BLASTP (top) or through genome mining (bottom). The cysimiditides expressed in this study are highlighted.

| **Organism** | ***O*-methyltransferase NCBI accession number** | **Precursor NCBI accession number (if given)** |
| --- | --- | --- |
| *Nocardiopsis sp. N85* | WP_274810681.1 | WP_274810680.1 |
| *Nocardiopsis sp. DSM 44743* | WP_311510783.1 | WP_274810680.1 |
| *Nocardiopsis dassonvillei* | WP_253792968.1 |  |
| *Nocardiopsis sp. FR26* | WP_160051417.1 | WP_160051416.1 |
| *Nocardiopsis sp. EMB25* | WP_268645093.1 | WP_268645092.1 |
| *Nocardiopsis sp. ATB16-24* | WP_285729426.1 | WP_285729427.1 |
| *Nocardiopsaceae bacterium* | MDA8370024.1 |  |
| *Salinactinospora qingdaonensis* | GAA3731431.1 |  |
| *Thermobifida cellulosilytica* | WP_083948071.1 |  |
| *Thermobifida alba* | WP_248591944.1 |  |
| *Microbispora sp.* | MBX6383762.1 |  |
| *Nocardiopsis sp. CC223A* | WP_306368270.1 | WP_306368271.1 |
| *Nocardiopsis sp. DSM 44743* | WP_311514221.1 | WP_311514222.1 |
| *Thermobifida fusca* | QOS59465.1  WP_227480949.1 |  |
| *Nocardiopsis sp. CT-R113* | WP_330095056.1 |  |
| *Nocardiopsis terrae* | WP_191275469.1 |  |
| *Nocardiopsis ganjiahuensis* | WP_017585468.1 |  |
| *Nocardiopsis sp. CNR-923* | WP_262391567.1 | WP_143831899.1 |
| *Nocardiopsis sp. B62* | WP_247666226.1 | WP_210843780.1 |
| *Nocardiopsis sp. SBT366* | WP_234004222.1 |  |
| *Spirillospora sp. NBC_01491* | WP_329521977.1 |  |
| *Actinomadura rugatobispora* | BFE30814.1 |  |
| *Streptomonospora sp. DSM 45055* | WP_311546033.1 |  |
| *Streptomonospora salina* | WP_246463618.1 |  |
| *Streptomonospora nanhaiensis* | WP_179767597.1 |  |
| *Streptomonospora nanhaiensis* | WP_217781190.1 |  |
| *Streptomonospora mangrovi* | WP_270072086.1 |  |
| *Nocardiopsis sp. CT-R113* | WP_330095078.1 |  |
| *Nocardiopsis deserti* | WP_223839422.1 |  |
| *Nocardiopsis dassonvillei* | WP_248770736.1 | WP_268742122.1 |
| *Nocardiopsis alborubida* | WP_061081300.1 | WP_267909101.1 |
| *Nocardiopsis alba* | WP_234305832.1 |  |
| *Nocardiopsis halotolerans* | WP_017569824.1 |  |
| *Nocardiopsis listeri* | WP_223809240.1 |  |
| *Nocardiopsis sp. ATB16-24* | WP_285730431.1 |  |
| *Nocardiopsis alkaliphila* | WP_017604998.1 |  |
| *Salinactinospora qingdaonensis* | GAA3754531.1 |  |
| --- | --- |  |
| *Streptomyces xiamenensis* | WP_046724513.1 |  |
| *Streptomyces sp. SBT349* | WP_049573986.1 |  |
| *Candidatus Frankia californiensis* | SBW17366.1 |  |
| *Frankia symbiont of Coriaria ruscifolia* | WP_131784212.1 |  |
| *Actinomadura syzygii* | WP_148347625.1 |  |
| *Frankia sp. CcI156* | ONH22856.1 |  |
| *Frankia sp. CgIM4* | OFB40277.1 |  |
| *Frankia sp. CeD* | WP_035911853.1 |  |
| *Frankia casuarinae* | ABD12601.1 |  |
| *Frankia sp. CcI156* | WP_076805381.1 |  |
| *Frankia sp. CeD* | WP_035910883.1 |  |
| *Frankia sp. Allo2* | WP_035732782.1 |  |
| *Frankia sp. B2* | WP_134351078.1 |  |
| *Frankia sp. CgIM4* | WP_070130813.1 |  |
| *Allosalinactinospora lopnorensis* | WP_046470800.1 |  |
| *Nocardiopsis sp. FIRDI 009* | WP_116247647.1 |  |
| *Allosalinactinospora lopnorensis* | WP_236700615.1 |  |
| *Nocardiopsis chromatogenes* | WP_017622943.1 |  |
| *Nocardiopsis baichengensis* | WP_017559121.1 |  |
